# Glucose Metabolism Controls Oxidative Burst and Lipid Mediator Production in Neutrophils upon Microbial Challenge

**DOI:** 10.1101/2025.08.31.672068

**Authors:** Alexandra Freitag, Kerstin Günther, Miriam Campillo Prados, Sophia M. Hochrein, Werner Schmitz, Katrin Sinning, Gunther Zischinsky, Bert Klebl, Knut Ohlsen, Joachim Morschhäuser, Oliver Werz, Paul M. Jordan, Martin Vaeth

## Abstract

Neutrophils are frontline responders against bacterial and fungal pathogens, requiring rapid energy and biosynthetic precursors to mount effective antimicrobial responses. To meet these demands, they primarily rely on aerobic glycolysis, making glucose uptake essential. Murine and human neutrophils express the glucose transporters GLUT1 and GLUT3; however, their specific roles in neutrophil immunobiology have not yet been fully elucidated. Here, we show that neutrophilic immune responses to *Candida albicans* and *Staphylococcus aureus* critically depend on GLUT1/3-dependent glucose uptake and glycolysis. Combined deletion of GLUT1 and GLUT3 almost completely abolished glucose uptake and aerobic glycolysis in murine neutrophils, yet did not impair granulopoiesis, indicating that homeostatic neutrophil development is largely independent of extracellular glucose. By contrast, during microbial challenge, loss of GLUT1/3 severely compromised NADPH-dependent ROS production, oxidative burst and cyclooxygenase-derived lipid mediator (LM) biosynthesis, demonstrating that glucose uptake via GLUT1/3 controls inflammatory effector functions of neutrophils. Moreover, genetic and pharmacologic inhibition of GLUT1/3-mediated glucose utilization reprograms neutrophil metabolism and LM biosynthesis toward an immunomodulatory phenotype. These findings identify a conserved nutrient-sensing metabolic checkpoint that governs neutrophil reprogramming and highlight novel opportunities for therapeutic immunomodulation.

## INTRODUCTION

Neutrophils are the most abundant leukocytes in human circulation and serve as a first-line defense against microbial invasion (Sender et al., 2023, Burn et al., 2021). As terminally differentiated granulocytes, neutrophils are rapidly recruited to sites of infection or damaged tissue, where they perform a broad range of antimicrobial, inflammatory and immunomodulatory functions (Evrard et al., 2018, Zhang et al., 2024). These functions include the phagocytosis of microbes, production of reactive oxygen species (ROS), degranulation, secretion of cytokines and chemokines, and the production of lipid mediators (LMs), as well as the formation of neutrophil extracellular traps (NETs) to eliminate invading microbes and prevent pathogen spreading (Burn et al., 2021). In addition to their role in antimicrobial defense, neutrophils orchestrate innate and adaptive immune responses through direct and indirect interaction with T cells, antigen-presenting cells and phagocytes, thereby shaping the local immune environment (Amulic et al., 2012). Neutrophils also play crucial roles in resolving inflammation, promoting tissue repair and restoring immune homeostasis by producing immunoregulatory molecules, releasing extracellular vesicles that dampen inflammation and facilitating the clearance of cellular debris (Bui et al., 2018, Kolaczkowska and Kubes, 2013, Ramos and Oehler, 2024). Although neutrophils are traditionally associated with a short lifespan and limited metabolic flexibility compared to other immune cells, they exhibit remarkable functional plasticity in response to environmental cues and pathogen-derived stimuli (Lahoz-Beneytez et al., 2016, Rogers and DeBerardinis, 2021).

The antimicrobial functions of neutrophils involve the generation of ROS to eliminate phagocytosed pathogens and the secretion of LMs, as potent signaling molecules with pleiotropic roles in immunity and inflammation (Ghodsi et al., 2024). Eicosanoids, such as leukotrienes (LTs) and prostaglandins (PGs), are well-characterized LMs that play crucial roles in initiating and amplifying inflammation by increasing vascular permeability, promoting leukocyte recruitment and contributing to coagulation (Dennis and Norris, 2015). In contrast, specialized pro-resolving mediators (SPMs) – including resolvins, protectins and maresins – represent a bovel class of LMs that actively promote the resolution of inflammation and support tissue repair (Jordan and Werz, 2022). LMs originate from membrane-derived polyunsaturated fatty acids (PUFAs), particularly arachidonic acid (AA), docosahexaenoic acid (DHA) and eicosapentaenoic acid (EPA), through tightly regulated enzymatic cascades. Cyclooxygenases (COX), lipoxygenases (LOX) and cytochrome P450 monooxygenases catalyze the conversion of PUFAs into a variety of bioactive LMs that collectively exert both pro-inflammatory and pro-resolving effects (Ghodsi et al., 2024, Jordan and Werz, 2022, Romp et al., 2020, Chandel, 2021). The spatiotemporal LM profile typically begins with pro-inflammatory PGs and LTs, which promote leukocyte recruitment to the site of inflammation. The release of LMs is tightly associated with a metabolic shift in neutrophils toward aerobic glycolysis to provide energy and enzymatic cofactors for the biosynthesis of pro-inflammatory eicosanoids (Zhang et al., 2024, Golenkina et al., 2024, Michaeloudes et al., 2020). As inflammation progresses, the LM profile undergoes a “class switch” toward pro-resolving lipids, such as SPMs and lipoxins, which terminate inflammation, facilitate tissue repair and restore immune homeostasis (Serhan, 2014, Albuquerque-Souza and Dalli, 2025).

Despite growing interest in immunometabolism, the interplay between neutrophil metabolism and LM secretion remains incompletely understood. Neutrophils have traditionally been considered primarily glycolytic; however, accumulating evidence suggests that oxidative phosphorylation (OXPHOS) and fatty acid metabolism also contribute to their differentiation and effector function (Jeon et al., 2020, Cao et al., 2022, Ettel and Weichhart, 2024, Lika et al., 2025). Most studies on the metabolic regulation of LM biosynthesis have focused on macrophages, where metabolic rewiring has been more extensively characterized (Giannakis et al., 2019, Günther et al., 2024, Werz et al., 2018, Jordan et al., 2020, Jordan et al., 2024). However, the specific contribution of central carbon metabolism – including glycolysis, the tricarboxylic acid (TCA) cycle and the pentose phosphate pathways (PPP) – to LM production has not yet been directly addressed. Crosstalk between glucose metabolism and LM biosynthesis is particularly relevant during antimicrobial responses, when neutrophils must rapidly mobilize energy reserves and provide metabolic intermediates to sustain effector functions. Several steps in LM metabolism require ATP and NADPH, which are closely linked to glycolysis, the PPP and the cellular redox balance (Smyrniotis et al., 2014). However, the precise contribution of glucose utilization and glycolytic metabolism to host defense and LM biosynthesis remains elusive.

In this study, we show that glucose uptake via the hexose transporters GLUT1 and GLUT3 controls antimicrobial effector function and LM production in both murine and human neutrophils. To address this, we employed two clinically relevant microorganisms from different kingdoms: *Candida albicans* (*C. albicans*), as an opportunistic fungal pathogen associated with mucosal and systemic infections in immunocompromised patients (Parambath et al., 2024), and *Staphylococcus aureus* (*S. aureus*), a gram-positive bacterium responsible for a broad spectrum of infections, ranging from soft tissue infections to life-threatening bacteremia (Howden et al., 2023). Using murine neutrophils with genetic ablation of GLUT1 and GLUT3, alongside pharmacological inhibition in human neutrophils, we investigated the role of GLUT1/3-mediated glucose uptake in responses to *C. albicans* and *S. aureus*. Through metabolic flux analysis, metabololipidomics and functional assays, we demonstrate that loss of GLUT1 and GLUT3 almost completely abolishes glucose uptake and aerobic glycolysis, resulting in a defective oxidative burst and a metabolic shift toward immunomodulatory pathways. Moreover, glucose utilization via GLUT1/3 governs the biosynthesis of LMs: suppression of glucose uptake shifts the LM profile from COX-derived PGs toward immunomodulatory species and enhances the secretion of the 5-LOX-derived chemotactic leukotriene LTB_4_. Notably, this glucose-dependent immunometabolic program is largely conserved in human neutrophils, identifying glucose uptake as a potential druggable metabolic checkpoint to modulate neutrophil effector functions and LM production in infectious and inflammatory diseases.

## RESULTS

### Neutrophils increase the glycolytic activity upon bacterial and fungal challenge

Neutrophils predominantly rely on aerobic glycolysis to meet their energetic and functional requirements, particularly during activation and microbial defense. Upon pathogen encounter, neutrophils rapidly increase their glucose uptake and glycolytic activity to support antimicrobial processes, including phagocytosis, ROS production and intracellular pathogen killing (Yipeng et al., 2024). Stimulation of murine *wild type* (WT) neutrophils with the fungal pathogen *Candida albicans* (strain SC5314) or the bacterial pathobiont *Staphylococcus aureus* (strain HG001) significantly increased the uptake of tritiated 2-desoxy-glucose ([^3^H]-2-DG) (**Fig. 1A**). Glucose serves as the universal fuel to immune cells and is imported from the extracellular environment via three structurally distinct transporter families. Sodium-glucose linked transporters (SGLTs) utilize extracellular sodium gradients to import glucose, while members of the SLC2A family (GLUTs) and the recently identified SLC50A1 (SWEET) proteins promote passive transport of hexoses across cellular membranes via facilitated diffusion. The GLUT family in humans comprises 14 members (GLUT1-14), of which 12 are conserved in mice (Deng and Yan, 2016, Mueckler and Thorens, 2013). Neutrophils predominantly express GLUT1 (*Slc2a1*) and GLUT3 (*Slc2a3*), while other members of the SLC2A family – as well as the SGLT and SWEET transporters – appear to be absent in granulocytes (Li et al., 2022, Wright, 2013, Watts et al., 2021). In resting murine neutrophils, the ‘neutron-type’ glucose transporter GLUT3 (*Slc2a3*) was predominantly expressed, whereas GLUT1 (*Slc2a1*) transcripts were detected at lower levels (**Fig. 1B** and **1C**). However, upon stimulation with *C. albicans* or *S. aureus*, the expression of *Slc2a1* was rapidly upregulated, while *Slc2a3* transcripts were repressed (**Fig. 1B** and **1C**). The shift in the expression of the two glucose transporters was also evident at the protein level, where GLUT1 was significantly increased in neutrophils after exposure to fungal and bacterial antigens (**Fig. 1D**). Although pathogen stimulation induces GLUT1 expression, the contribution of the two hexose transporters to glucose uptake, metabolic reprogramming and the impact on neutrophil effector functions remains incompletely understood.

**Figure 1.**
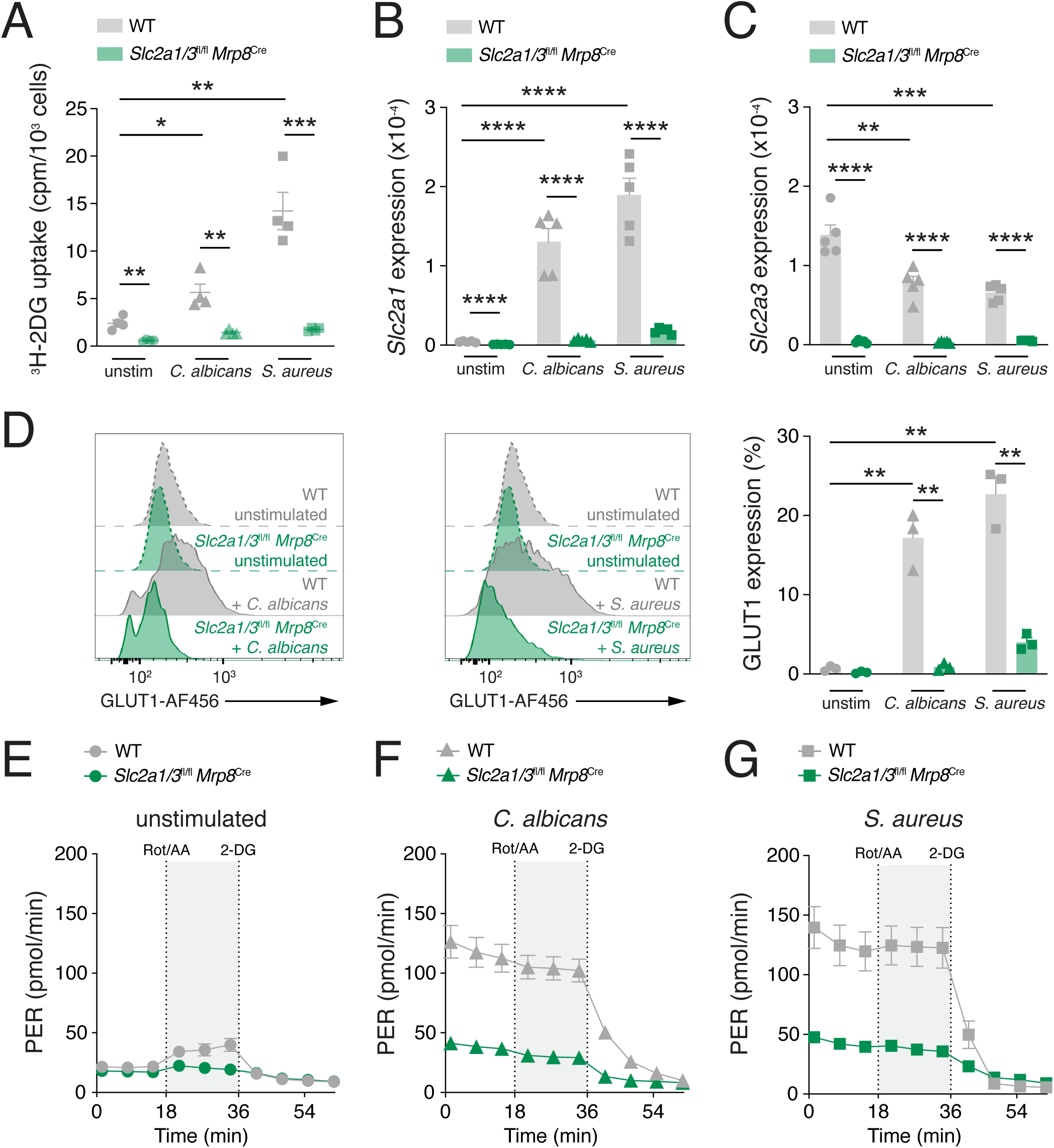
Neutrophils increase their glycolytic flux upon *C. albicans* and *S. aureus* stimulation. (**A**) Analysis of tritiated 2-dexoxy-glucose ([^3^H]-2-DG) uptake by WT and GLUT1/3-deficient neutrophils stimulated for 6 h with inactivated *C. albicans* (hyphae) or *S. aureus* (both at MOI = 5) using a scintillation counter; means ± SEM of 4 biological replicates per group. (**B** and **C**) Analysis of *Slc2a1* (GLUT1) **(B)** and *Slc2a3* (GLUT3) **(C)** gene expression in bone marrow (BM) neutrophils from WT and GLUT1/3-deficient (*Slc2a1*^fl/fl^ *Slc2a3*^fl/fl^ *Mrp8*^Cre^) mice stimulated for 6 h with live *C. albicans* (hyphae) or *S. aureus* (both at MOI = 1) by qRT-PCR; mean ± SEM of 5 biological replicates per group. **(D)** Flow cytometric analysis of GLUT1 protein expression of WT and GLUT1/GLUT3-deficient BM neutrophils stimulated for 6 h with live *C. albicans* (hyphae) or *S. aureus* (both at MOI = 0.4); mean ± SEM of 3 biological replicates per group. **(E-G)** Glycolytic proton efflux rate (PER) analyses of WT and GLUT1/3-deficient neutrophils. Cells were left unstimulated **(E)** or treated for 6 h with inactivated *C. albicans* (hyphae) **(F)** and *S. aureus* **(G)** (both at MOI = 5) using a Seahorse extracellular flux analyzer; mean ± SEM of 4 biological replicates per group. *p < 0.05, **p < 0.01, ***p < 0.001, ****p < 0.0001 by unpaired Student’s t test (A-D)

### Ablation of GLUT1 and GLUT3 abolishes glucose uptake and glycolysis in neutrophils

To investigate the roles of GLUT1 and GLUT3 in neutrophil development, metabolism and function, we generated conditional knockout mice by crossing *Slc2a1*^fl/fl^ (Macintyre et al., 2014) and *Slc2a3*^fl/f^ (Fidler et al., 2017) mice with *Mrp8*^Cre^ transgenic animals (Passegué et al., 2004). The *Mrp8* gene (encoding S100A8) is typically not expressed in granulocyte-monocyte progenitors (GMPs) but upregulated in neutrophil progenitors (preNeu) and progressively expressed in immature and mature neutrophil subsets. Importantly, *Mrp8*-driven Cre recombinase is not expressed in other myeloid cell types, and preNeu cells exclusively give rise to the neutrophil lineage (Evrard et al., 2018). Thus, the resulting *Slc2a1/3*^fl/fl^*Mrp8*^Cre^ mice lacked both GLUT1 and GLUT3 specifically in neutrophils (**Fig. 1B** and **C**), providing a suitable model to investigate the role of GLUT1/3-mediated glucose uptake in neutrophil immune responses. *Slc2a1/3*^fl/fl^*Mrp8*^Cre^ mice were born at expected Mendelian ratios and did not exhibit any overt immune phenotype compared to their WT littermate controls. Intriguingly, combined deletion of GLUT1 and GLUT3 did not perturb homeostatic granulopoiesis. Neutrophil frequencies and absolute numbers across maturation states remained comparable between WT and *Slc2a1/3*^fl/fl^*Mrp8*^Cre^ mice in the BM, blood and peripheral lymphoid organs (**Fig. S1A-C**).

The expression of GLUT1 and GLUT3 at both mRNA and protein levels was undetectable in resting and pathogen-stimulated neutrophils from *Slc2a1/3*^fl/fl^*Mrp8*^Cre^ mice (**Fig. 1B-D**), confirming efficient deletion of both transporters. Consistently, uptake of tritiated 2-DG (**Fig. 1A**) and lactate-dependent extracellular acidification – measured as the proton efflux rate (PER) using a Seahorse extracellular flux analyzer (F. Gubert et al., 2025) – were almost completely abolished in both resting and pathogen stimulated GLUT1/3-deficient neutrophils (**Fig. 1E-G**), indicating that the loss of GLUT1 and GLUT3 is not compensated by other hexose transporters. In unstimulated WT neutrophils, basal and maximal glycolysis was relatively low (**Fig. 1E**), whereas lactate secretion (measures as PER) was markedly increased upon stimulation with inactivated *C. albicans* or *S. aureus* (**Fig. 1F** and **1G**). By contrast, GLUT1/3-deficient neutrophils failed to upregulate their glycolytic activity, which remained at basal levels following microbial stimulation (**Fig. 1F** and **1G**). Moreover, no compensatory increase in mitochondrial respiration – measured as oxygen consumption rate (OCR) – was observed in GLUT1/3-deficient neutrophils compared to WT controls (**Fig. S2A-C**). Thus, pathogen-stimulated WT neutrophils adopt a glycolytic metabolic profile without a corresponding increase in mitochondrial respiration, whereas GLUT1/3-deficient neutrophils remain in a metabolically quiescent state (**Fig. S2D**). Notably, activation or priming of GLUT1/3-deficient neutrophils was comparable to that of WT controls following stimulation with *C. albicans* or *S. aureus*, as evidenced by CD62L shedding (**Fig. S2E**) and the upregulation of CD11b (**Fig. S2F**) (Ivetic et al., 2019).

These findings suggest that glucose uptake and glycolytic metabolism in pathogen-stimulated murine neutrophils are predominantly mediated by GLUT1 and GLUT3. Intriguingly, both glucose transporters seem dispensable for homeostatic granulopoiesis and neutrophil homing to secondary lymphoid organs under steady-state conditions.

### Extracellular glucose uptake fuels the pentose phosphate pathways and ROS production

To elucidate the metabolic consequences of GLUT1/3 ablation in greater detail, we performed stable isotope tracing of polar metabolites using ^13^C-labelled glucose, analyzed by mass spectrometry coupled with liquid chromatography (LC-MS). To avoid confounding effects from pathogen-derived metabolites – which are abundant in live and inactivated fungi and bacteria and can be extracted by phagocytes to fuel metabolic functions (Lesbats et al., 2025, Wise et al., 2025) – we employed the insoluble fungal β-1,3-glucan zymosan as a model stimulus. This strategy enabled us to mimic fungal infection while minimizing interference in our LC–MS analyses caused by pathogen-derived metabolites and metabolic competition between pathogens and neutrophils.

WT and GLUT1/3-deficient neutrophils were stimulated with zymosan for 12 hours in medium containing [U-^13^C_6_]-glucose. Following stimulation, cells were washed, and the pellets were analyzed with LC-MS. Polar metabolites derived from ^13^C_-_glucose were identified based on their isotopic mass shifts – for example, three-carbon metabolites such as pyruvate or lactate exhibited an m+3 isotopic signature (**Fig. 2A**). Interestingly, even in the absence of stimulation, global metabolic differences were observed between WT and GLUT1/3-deficient neutrophils, affecting TCA cycle intermediates, as well as purine, pyrimidine and glycolytic metabolites (**Fig. 2B**). Zymosan stimulation further amplified these differences (**Fig. 2C**). Notably, several key metabolites of the glycolytic pathway, including the end products pyruvate and lactate were among the most significantly downregulated metabolites in GLUT1/3-deficient neutrophils compared to WT controls (**Fig. 2C** and **2D**), consistent with their impaired glycolytic flux observed in Seahorse analyses (**Fig. 1E-G**). Beyond glycolysis, glucose can be diverted into the pentose phosphate pathway (PPP) to support nucleotide biosynthesis and generate nicotinamide adenine dinucleotide phosphate (NADPH) – a key reducing equivalent for anabolic processes and cellular redox regulation (O’Neill et al., 2016). In the oxidative branch of the PPP, glucose-6-phosphate (G6P) is converted to ribulose-5-phosphate, yielding NADPH and CO_2_. Alternatively, glucose can be directly oxidized to gluconate with concomitant NADPH production and subsequently phosphorylated to 6-phospho-gluconate (TeSlaa et al., 2023). Ribulose-5-phosphate enters the non-oxidative phase of the PPP, where it is metabolized into various sugars, including ribose-5-phosphate – an essential precursor for the purine and pyrimidine biosynthesis – or it can cycle back into glycolysis or the PPP itself (**Fig. 2A**). Activated neutrophils engage the oxidative branch of the PPP to generate NADPH, which fuels ROS production via the NADPH-oxidase complex – critical for intracellular pathogen killing and the formation of neutrophil extracellular traps (NETs) (Britt et al., 2022, Azevedo et al., 2015). In addition to the glycolytic end products pyruvate and lactate, we observed a marked increase of ^13^C-labelled gluconate in WT neutrophils following zymosan stimulation, which was nearly absent in GLUT1/3-deficient neutrophils (**Fig. 2D**). Intriguingly, glucose-derived *de novo* synthesis of purine nucleotides and nucleosides, (AMP, adenosine, inosine and xanthine) was not impaired. In fact, GLUT1/3-deficient neutrophils showed higher levels of ^13^C-labelled purine metabolites after zymosan stimulation (**Fig. 2E**), indicating an imbalance between the oxidative and non-oxidative branches of the PPP. The abundance of ^13^C-labelled intermediates of the TCA cycle, including citrate and aconitate, was modestly increased in GLUT1/3-deficient neutrophils compared to WT controls following zymosan stimulation (**Fig. 2F**), suggesting that mitochondrial metabolism remains functionally intact. These findings align with the unchanged oxygen consumption rates observed in GLUT1/3-deficient neutrophils (**Fig. S2A-C**). Notably, the production of the anti-inflammatory TCA cycle metabolite itaconate was significantly elevated in GLUT1/3-deficient neutrophils following zymosan stimulation (**Fig. 2F**), suggesting a metabolic shift toward immunomodulatory pathways when extracellular glucose utilization is impaired.

**Figure 2.**
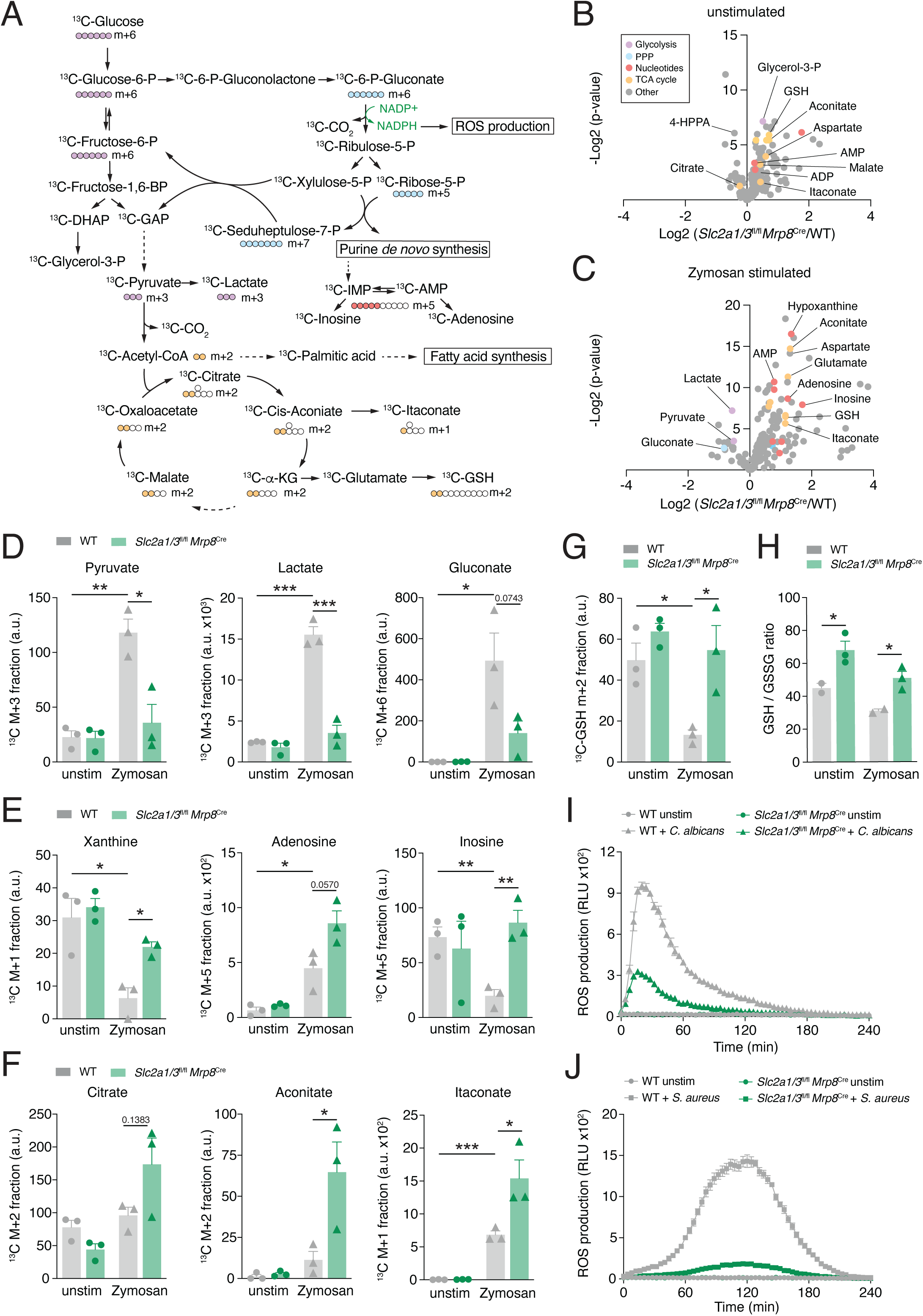
Metabolic profiling of GLUT1/3-deficient neutrophils. **(A)** Schematic representation of stable isotope tracing of ^13^C-glucose in WT and GLUT1/3-deficient neutrophils using high-resolution mass spectrometry coupled to liquid chromatography (LC-MS). (**B** and **C**) Volcano plots of differential metabolite levels between WT and GLUT1/3-deficient neutrophils without stimulation **(B)** and after Zymosan **(C)** stimulation for 12 h; mean ± SEM of 3 biological replicates per group. **(D-G)** Analysis of selected isotope-labelled polar metabolites derived from ^13^C-glucose in WT and GLUT1/3-deficient neutrophils; mean ± SEM of 3 biological replicates per group. **(H)** Ratio of total (unlabeled) GSH to GSSG in WT and GLUT1/3-deficient neutrophils; mean ± SEM of 3 biological replicates per group. (**I** and **J)** Reactive oxygen species (ROS) production in WT and GLUT1/3-deficient neutrophils stimulated for with live *C. albicans* (hyphae) **(I)** or *S. aureus* **(J)** (both at MOI = 1) over time using a luminol-based ROS assay; mean ± SEM of 5 biological replicates per group. *p < 0.05, **p < 0.01, ***p < 0.001 by unpaired Student’s t test (D-H)

The respiratory burst, driven by the PPP-fueled NADPH oxidase complex, is essential for intracellular pathogen killing and the formation of NETs (Britt et al., 2022). To manage the oxidative stress generated by this burst, neutrophils rely on the glutathione (GSH) antioxidant system to regulate ROS levels (Britt et al., 2022, Forman et al., 2009). In WT neutrophils, GSH synthesis was markedly reduced upon zymosan stimulation, thereby enabling elevated ROS production. In contrast, GLUT1/3-deficient neutrophils failed to downregulate GSH biosynthesis (**Fig. 2G**), suggesting that they do not initiate a respiratory burst reaction under these conditions. The cellular redox status, reflected by the ratio of reduced (GSH) to oxidized (GSSG) glutathione, was also significantly higher in GLUT1/3-deficient neutrophils compared to WT controls (**Fig. 2H**). These elevated GSH/GSSG ratios indicate a lower level of redox stress, further supporting the notion that an oxidative burst does not occur in the absence of GLUT1 and GLUT3. Correspondingly, GLUT1/3-deficient neutrophils failed to produce ROS in response to *C. albicans* (**Fig. 2I**) or *S. aureus* (**Fig. 2J**) stimulation.

Collectively, neutrophils rely on extracellular glucose uptake via GLUT1 and GLUT3 to fuel glycolysis and NADPH production through the oxidative branch of the PPP. In absence of GLUT1/3-mediated glucose metabolism, neutrophils fail to mount an efficient NADPH-dependent respiratory burst upon pathogen stimulation and exhibit a metabolic shift toward immunomodulatory pathways.

### GLUT1/3-mediated glucose utilization regulates the secretion of lipid mediators

Although the NADPH oxidase complex is the primary source of superoxide during the oxidative burst in phagocytes, additional metabolic pathways also contribute to ROS generation and redox regulation. Beyond mitochondrial respiration, the oxidative metabolism of AA via cyclooxygenases (COX) and lipoxygenases (LOX) may further support cellular ROS production in neutrophils and macrophages (Zhao et al., 2013, Hu et al., 2017). Elevated ROS levels can, in turn, enhance the expression and activity of COX and LOX enzymes, establishing a feed-forward loop that not only amplifies antimicrobial effector functions but also promotes the production of immunomodulatory LMs, such as PGs, LTs and SPMs (Golenkina et al., 2024) (**Fig. 3A**). These LMs play critical roles in antimicrobial immunity, orchestrate broader immune responses and contribute to the resolution of inflammation (Brennan et al., 2021, Shimizu, 2009).

**Figure 3.**
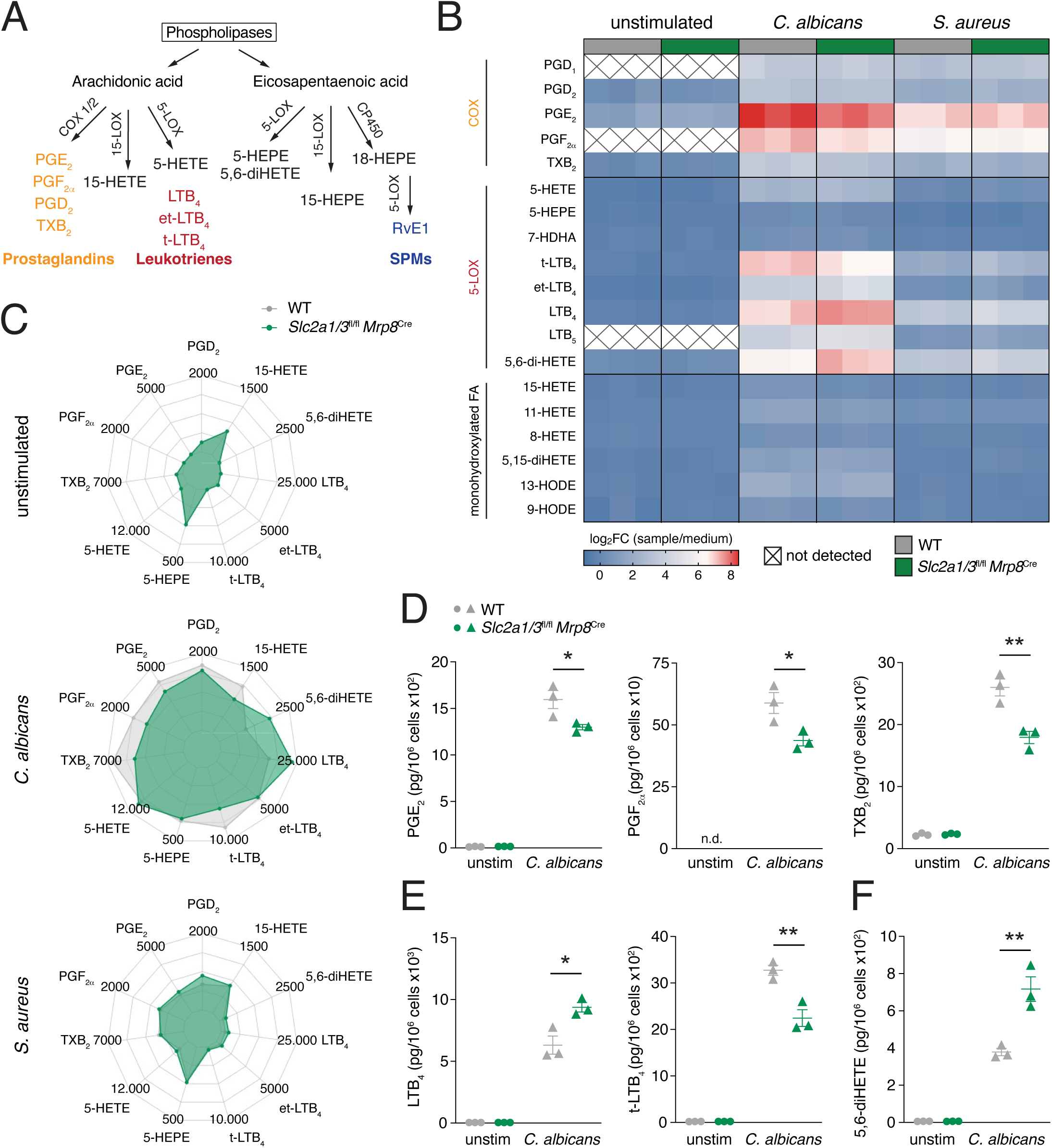
Ablation of GLUT1/3 impacts the secretion of inflammatory lipid mediators. **(A)** Scheme of lipid mediator biosynthesis by COX and LOX-dependent pathways. **(B)** Heatmap of lipid mediator (LM) levels in the supernatant of WT and GLUT1/3-deficient (*Slc2a1*^fl/fl^ *Slc2a3*^fl/fl^ *Mrp8*^Cre^) neutrophils stimulated for 6 h with live *C. albicans* (hyphae) or *S. aureus* (both at MOI = 1) using ultra performance liquid chromatography – tandem and mass spectrometry (UPLC-MS/MS). Heatmaps show Log_2_-transformed fold changes relative to medium control, with values ranging from low (blue) to high (red); Each condition includes 3 biological replicates per group. (**C** and **D**) Radar plots (**C**) and bar graphs (**D**) of lipid mediator amounts in the supernatant of WT and GLUT1/GLUT3-deficient neutrophils left unstimulated or stimulated for 6 h with live *C. albicans* (hyphae) or *S. aureus* (both at MOI = 1). Data shown as pg/2.5x10^6^ cells in (C) and pg/10^6^ cells in (D); mean ± SEM of 3 biological replicates per group. *p < 0.05, **p < 0.01, ***p < 0.001 by unpaired Student’s t test (D)

The two COX isoforms, COX-1 and COX-2, are membrane-bound enzymes that catalyze the conversion of AA into the intermediate PGH_2_. Subsequently, specific prostaglandin and thromboxane synthases convert PGH_2_ into prostanoids, including PGE_2_, PGF_2α_, PGD_2_, and the thromboxanes TXA_2_ and TXB_2_ (Smith et al., 2011, Shimizu, 2009) (**Fig. 3A**). Besides COX enzymes, the enzyme 5-lipoxygenase (5-LOX) initiates the biosynthesis of LTs, potent mediators of the inflammatory response. This occurs through a two-step reaction that starts with the oxygenation of the substrate AA to generate the intermediate 5-S-hydroperoxyeicosatetraenoic acid (5-S-HpETE). A second hydrogen abstraction converts 5-S-HpETE into LTA_4_, which is then metabolized to the potent leukocyte chemoattractant LTB_4_ via LTA_4_ hydrolase, or conjugated with GSH to generate the cysteinyl-LTs C_4_, D_4_, and E_4_ that mediate bronchoconstriction and increase vascular permeability (Shimizu, 2009) (**Fig. 3A**). In addition to COX and LOX enzymes, cytochrome P450 (CYP450) enzymes also metabolize PUFAs into diverse bioactive LMs, including hydroxyeicosatetraenoic acids (HETEs), hydroxyeicosapentaenic acids (HEPEs), epoxyeicosatrienoic acids (EETs) and hydroxyoctadecadienoic acids (HODEs) (**Fig. 3A**).

To assess how glucose metabolism affects LM production, we performed metabololipidomics with supernatants from WT and GLUT1/3-deficient neutrophils stimulated with *C. albicans* or *S. aureus* using ultra-performance liquid chromatography coupled with tandem mass spectrometry (UPLC-MS/MS) (**Fig. 3B-G** and **Fig. S3**). In the absence of stimulation, both WT or GLUT1/3-deficient neutrophils released only low or undetectable levels of PGs and LTs. Upon stimulation, *C. albicans* elicited a robust production and secretion of a broad range of LMs, while *S. aureus* induced only a modest increase in bioactive LM levels (**Fig. 3B** and **3C**, **Table S1**). Although *S. aureus* stimulation showed minimal differences between WT and GLUT1/3-deficient neutrophils, challenge with *C. albicans* led to pronounced differences in multiple LM species (**Fig. 3B**, **3C** and **Fig. S3**). Among the affected LMs, levels of PGE_2_, PGF_2α_, TXB_2_, t-LTB_4_, monohydroxylated fatty acids (8-HETE, 11-HETE, 15-HETE), as well as 9-HODE, were significantly reduced, whereas LTB_4_ and 5S,6R-diHETE were markedly increased in GLUT1/3-deficient neutrophils compared to WT controls (**Fig. 3B-F**). These results suggest that COX-dependent production of PGs – particularly PGE_2_, PGF_2α_ and TXB_2_ – depends on GLUT1/3-mediated glucose uptake during *C. albicans* stimulation. In contrast, 5-LOX and CYP450-derived products appears more complex. While LTB_4_ and LTB_5_ were significantly increased in GLUT1/3-deficient neutrophils, t-LTB_4_ and et-LTB_4_ – non-enzymatic hydrolysis products of LTA_4_ (Haeggstrom, 2000) – were reduced (**Fig. 3E**). These findings indicate that glucose metabolism differentially modulates LOX enzymes activity during fungal stimulation. Notably, the production of the immunoregulatory lipid 5S,6R-diHETE was markedly enhanced in GLUT1/3-deficient neutrophils following *C. albicans* exposure (**Fig. 3F**). 5S,6R-diHETE possesses anti-inflammatory and tissue-protective properties (Kobayashi et al., 2021), supporting the notion that glucose restriction shifts the LM profile from COX toward LOX-mediated pathways in neutrophils.

Taken together, while loss of glucose uptake does not fully abrogate LM secretion, GLUT1/3-deficent neutrophils display a marked reduction in COX-derived prostaglandins and altered production of LOX-derived LTB_4_ and monohydroxylated LMs following *C. albicans* stimulation.

### GLUT1/3-mediated glucose uptake and metabolism are conserved in human neutrophils

To determine whether our findings in GLUT1/3-deficient murine bone marrow neutrophils extend to human neutrophils, we examined glucose uptake and immunometabolism in human peripheral blood neutrophils (PBNs). Like murine neutrophils, human neutrophils exhibited low basal GLUT1 expression, whereas GLUT3 was robustly expressed even in unstimulated cells (**Fig. 4A** and **B**). Upon microbial challenge, GLUT1 was significantly upregulated by both *C. albicans* and *S. aureus* (**Fig. 4A**), mirroring murine responses. By contrast, GLUT3 expression increased in response to *C. albicans* and remained unchanged following *S. aureus* stimulation (**Fig. 4B**), suggesting that both transporters contribute to the immunometabolic response and effector function of human neutrophils.

**Figure 4.**
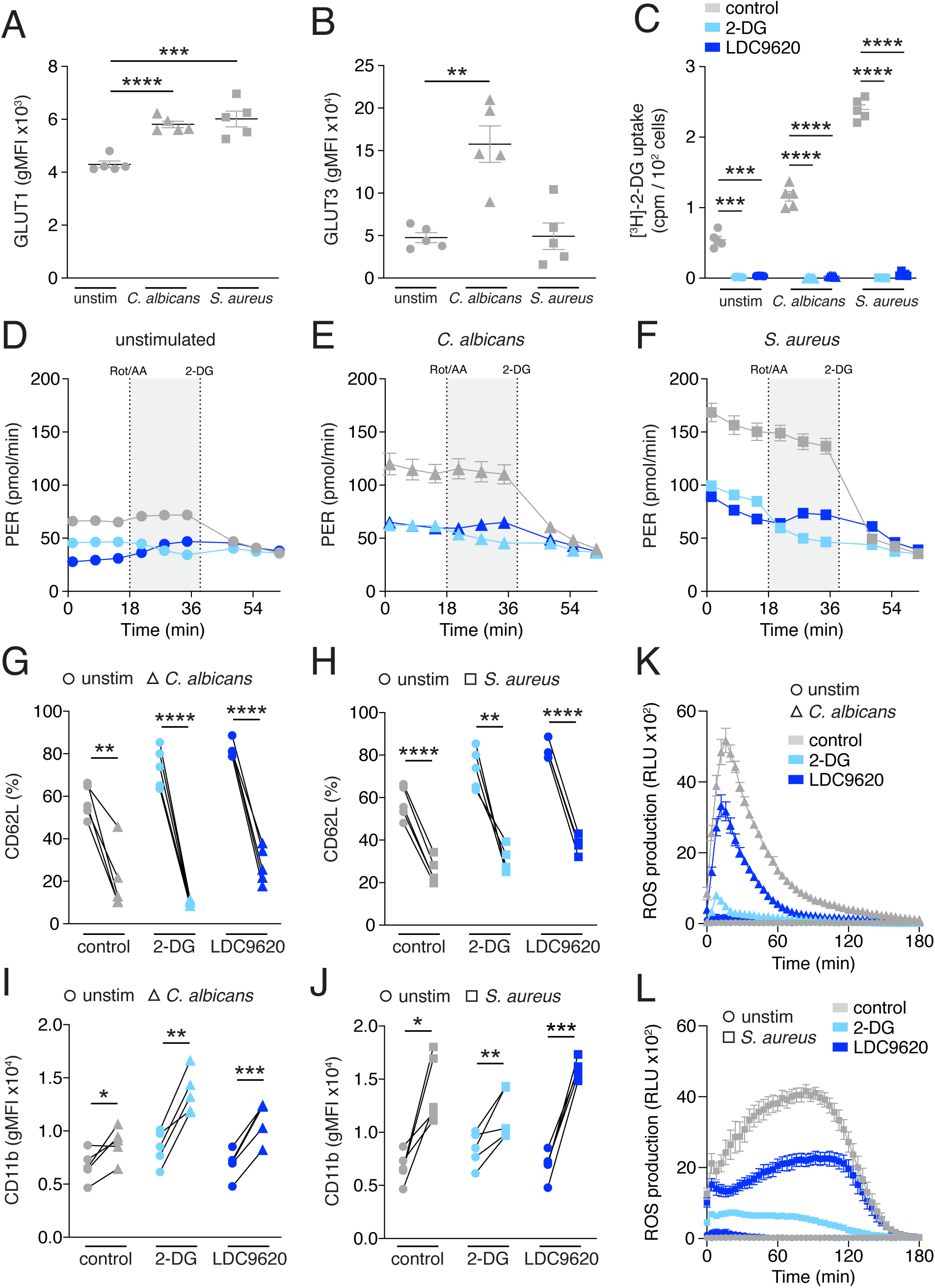
Inhibition of GLUT1/3-mediated glucose utilization by human neutrophils. (**A** and **B**) Analysis of *Slc2a1* (GLUT1) **(A)** and *Slc2a3* (GLUT3) **(B)** gene expression in human blood neutrophils stimulated for 3 h with live *C. albicans* (hyphae) or *S. aureus* (both at MOI = 1) using PrimeFlow; mean ± SEM of 5 donors. **(C)** Analysis of tritiated 2-desoxy-glucose ([^3^H]-2-DG) uptake by control, 2-DG and GLUT1/3-inhibitor (LDC9620) treated human neutrophils stimulated for 3 h with inactivated *C. albicans* (hyphae) or *S. aureus* (both at MOI = 5) using a scintillation counter; means ± SEM of 5 donors. (**D** – **F**) Analysis of glycolytic proton efflux rate (PER) in control, 2-DG and GLUT1/3-inhibitor treated human neutrophils left unstimulated (**D**) or stimulated with inactivated *C. albicans* (hyphae) (**E**) or *S. aureus* (**F**) for 3 h (both at MOI = 5) using Seahorse extracellular flux analyzer; mean ± SEM of 5 donors. (**G** and **H**) Analysis of CD62L expression of control, 2-DG and GLUT1/3-inhibitor treated human neutrophils left unstimulated or stimulated for 3 h with live *C. albicans* (hyphae) **(G)** or *S. aureus* **(H)** (both at MOI = 1); mean ± SEM of 5 donors. (**I** and **J**) Analysis of CD11b expression of control, 2-DG and GLUT1/3-inhibitor treated human neutrophils left unstimulated or stimulated for 3 h with live *C. albicans* (hyphae) **(I)** or *S. aureus* **(J)** (both at MOI = 1); mean ± SEM of 5 donors. (**K** and **L**) Reactive oxygen species (ROS) production by control, 2-DG and GLUT1/3-inhibitor treated human neutrophils stimulated with live *C. albicans* (hyphae) **(K)** and *S. aureus* **(L)** (both MOI = 1) using a luminol-based ROS assay; mean ± SEM of 5 donors. *p < 0.05, **p < 0.01, ***p < 0.001, ****p < 0.0001 by paired Student’s t test (A-C, G-J)

PBNs are terminally differentiated, short-lived neutrophils with a lifespan of ∼1-2 days and no capacity for proliferation, making genetic manipulation techniques, such as CRISPR/Cas9-mediated genome editing, ineffective. Their high sensitivity to stress during manipulation further limits genetic interventions. Consequently, pharmacological inhibition of glucose uptake and metabolism represents the most practical approach to investigate the role of GLUT1 and GLUT3 in human neutrophils. Thus, we employed two complementary pharmacological strategies to dissect the role of glucose metabolism in PBNs. First, we used 2-desoxy-glucose (2-DG), a structural glucose analogue that enters cells via glucose transporters. Intracellularly, 2-DG is rapidly phosphorylated to 2-DG-6-phosphate, which competitively inhibits hexokinases and other glycolytic enzymes, broadly suppressing glucose uptake and utilization (Pajak et al., 2019, Wu et al., 2023). In parallel, we applied a highly specific inhibitor of GLUT1 and GLUT3 (LDC9620), to directly assess GLUT1/3-dependent glucose uptake in human neutrophils (**Fig. 4C-L**).

Stimulation of PBNs with *C. albicans* or *S. aureus* significantly increased the uptake of the tritiated glucose analogue 2-DG (**Fig. 4C**) and enhanced their glycolytic activity, as indicated by elevated lactate-dependent extracellular acidification rates (**Fig. 4D-F**). Both 2-DG and LDC9620 abolished glucose uptake (**Fig. 4C**) and aerobic glycolysis (**Fig. 4D-F**), confirming their suitability as pharmacological tools to investigate GLUT1/3 function in human neutrophils. Notably, the GLUT1/3-specific inhibitor LDC9620 suppressed glucose uptake and glycolysis to a similar extent as 2-DG, indicating that the acute blockade of GLUT1 and GLUT3 in human neutrophils is also not compensated by other hexose transporters. Similar to the findings in GLUT1/3-deficient murine neutrophils, inhibition of glucose uptake did not impair PBN activation, as shown by preserved CD62L shedding (**Fig. 4G** and **4H**) and CD11b upregulation (**Fig. 4I** and **J**) following *C. albicans* or *S. aureus* stimulation. Given that GLUT1/3-deficient murine neutrophils exhibited severe defects in the oxidative burst in response to microbial challenge, we next examined whether PBNs display similar impairment after pharmacological GLUT1/3 inhibition. Strikingly, both 2-DG and LDC9620 markedly suppressed ROS production in response to *C. albicans* (**Fig. 4K**) and *S. aureus* (**Fig. 4L**), revealing that the roles of GLUT1 and GLUT3 in glucose uptake, glycolytic metabolism and oxidative responses are largely conserved between murine and human neutrophils.

### Inhibition of glucose utilization modulates lipid mediator production in human neutrophils

Having established that GLUT1 and GLUT3 play a similar role in glucose uptake and NADPH-dependent ROS production in human neutrophils, we next examined whether glucose metabolism also regulates the biosynthesis and secretion of LMs in PBNs. Using UPLC-MS/MS profiling, we observed that human neutrophils exhibited a LM signature similar to that of murine neutrophils upon stimulation with *C. albicans* or *S. aureus*. Unstimulated PBNs released only basal or undetectable amounts of bioactive lipids, whereas microbial challenge markedly enhanced their production of LMs, with *C. albicans* eliciting a stronger response than *S. aureus* (**Fig. 5A** and **5B**, **Table S2**).

**Figure 5.**
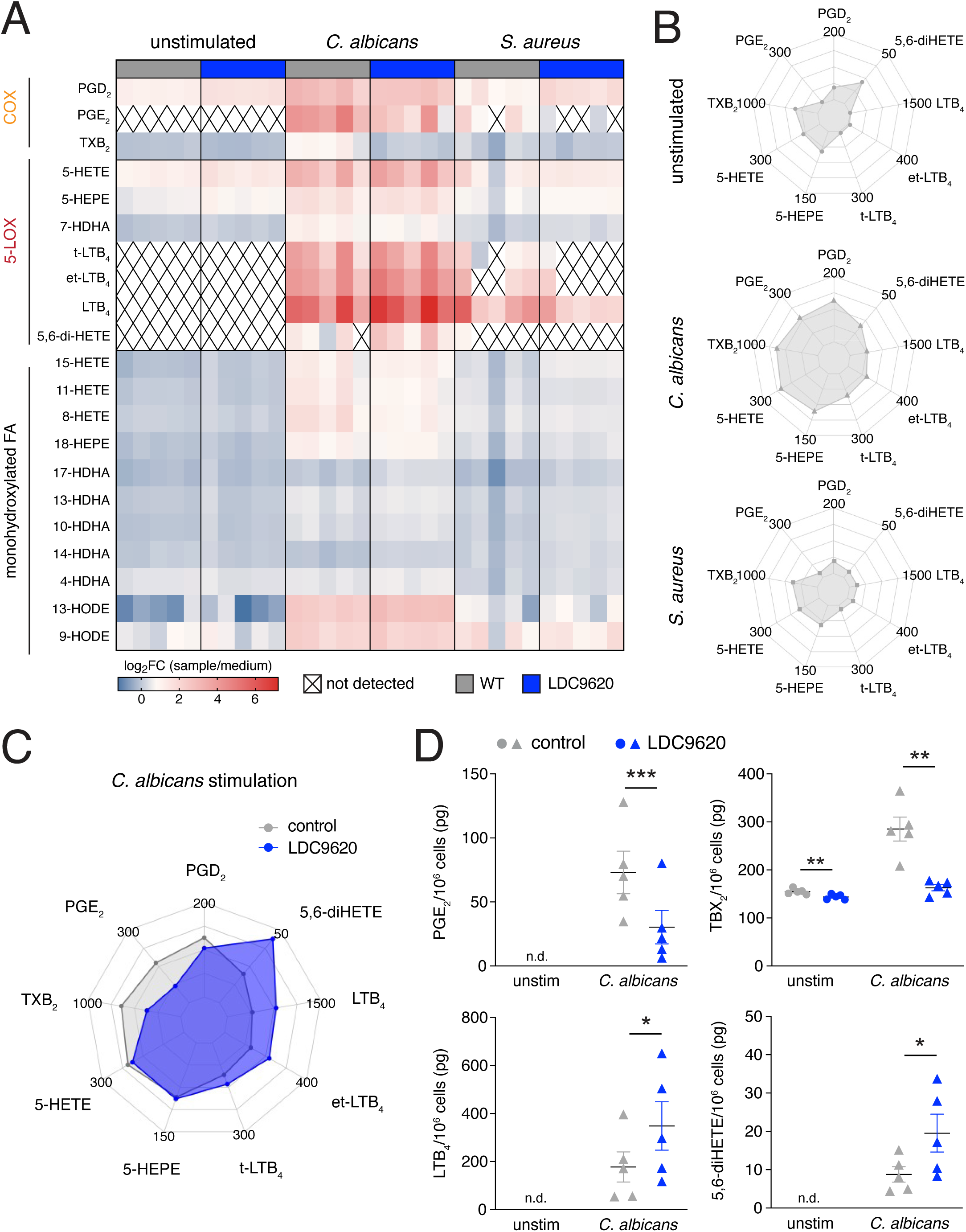
Acute inhibition of GLUT1/3 in human neutrophils alters lipid mediator production. **(A)** Heatmap of lipid mediators in the supernatants of control and GLUT1/3-inhibitor (LDC9620, 4 µM) treated human neutrophils stimulated with live *C. albicans* (hyphae) or *S. aureus* (both at MOI = 1) for 3 h using ultra performance liquid chromatography – tandem and mass spectrometry (UPLC-MS/MS). Heatmaps show Log_2_-transformed fold changes relative to medium control, with values ranging from low (blue) to high (red); Each condition includes 5 donors per group. **(B)** Radar plots of lipid mediator amounts in the supernatant from human neutrophils left unstimulated or stimulated with live *C. albicans* (hyphae) or *S. aureus* (both at MOI = 1) for 3 h. Data shown as pg/2.25x10^6^ cells. Mean from n = 5. **(C)** Radar plots of lipid mediator concentrations in the supernatant of control and GLUT1/3-inhibitor treated human neutrophils stimulated with live *C. albicans* (hyphae) (MOI = 1) for 3 h. Data given as pg/2.25x10^6^ cells. **(D)** Quantification of selected lipid mediators secreted from control and GLUT1/3-inhibitor treated human neutrophils left unstimulated or stimulated for 3 h with live *C. albicans* (hyphae) (MOI = 1). Data given as pg/10^6^ cells; mean ± SEM of 5 donors. *p < 0.05, **p < 0.01, ***p < 0.001 by paired Student’s t test (D); n.d. = not detected

Upon *C. albicans* stimulation, pharmacological inhibition of GLUT1/3 in human neutrophils led to a marked reduction in COX-derived PGs, including PGE_2_, PGD_2_ and TXB_2_ (**Fig. 5A-D**). In contrast, LOX and CYP450-dependent metabolites showed a more nuanced response: levels of certain monohydroxylated eicosatetraenoic acids – such as 8-HETE, 11-HETE and 15-HETE –decreased after GLUT1/3-blockade, whereas LTB_4_ (**Fig. 5C, 5D** and **Fig. S4**) and monohydroxylated docosahexaenoic acid (HDHA) species – including 4-HDHA, 10-HDHA, 13-HDHA, 14-HDHA and 17-HDHA – were significantly elevated (**Fig. 5A**). Notably, treatment with the GLUT1/3-specific inhibitor LDC9620 also enhanced the production of 5S,6R-diHETE in PBNs (**Fig. 5D**), indicating that glucose metabolism influences COX and LOX-mediated LM pathways in human neutrophils, similar to the observations in murine neutrophils.

Altogether, our study reveals that GLUT1/3-mediated glucose uptake is a conserved regulator of neutrophil effector functions and LM biosynthesis during fungal and bacterial challenges. Both genetic and pharmacological inhibition of glucose metabolism in murine and human neutrophils suppress NADPH-dependent respiratory burst and COX-derived PG production, while redirecting cellular metabolism toward LOX pathways. These findings establish GLUT1/3-mediated glucose uptake as a critical metabolic checkpoint with promising therapeutic potential for immunomodulation.

## DISCUSSION

Aulus Cornelius Celsus, a Roman encyclopedist, provided the first recorded definition of inflammation: “*Notae vero inflammationis sunt quatuor: rubor et tumor cum calore et dolore”*, identifying the four cardinal signs of acute inflammation as redness (*rubor*), swelling (*tumor*), heat (*calor*) and pain (*dolor*) (Cavaillon, 2021). Central to inflammatory response are neutrophils – rapidly recruited phagocytes that constitute the first line of defense against tissue injury and invading pathogens. Upon activation, neutrophils initiate and amplify key inflammatory processes that directly contribute to these cardinal signs, including vascular permeability, inflammation and the recruitment of leukocytes to the sites of inflammation. These effects are mediated through the release of ROS, pro-inflammatory cytokines and chemokines and the secretion of bioactive LMs (Burn et al., 2021). Importantly, neutrophils are not only drivers of inflammation but also play a crucial role in its resolution and the restoration of tissue homeostasis through mechanisms that include the clearance of pathogens and cellular debris, as well as the production of pro-resolving proteins and LMs (Serhan, 2014).

A defining feature of neutrophil activation is the rapid and profound metabolic shift toward aerobic glycolysis, which fuels antimicrobial and inflammatory functions (Li et al., 2022, Michaeloudes et al., 2020, Britt et al., 2022). In this study, we identify a critical role for the glucose transporters GLUT1 and GLUT3 in orchestrating the immunometabolic program of neutrophils upon exposure to bacterial and fungal pathogens. Using mice with neutrophil-specific deletion of *Slc2a1* and *Slc2a3* (encoding GLUT1 and GLUT3), together with pharmacological inhibition of glucose utilization in human neutrophils, we here demonstrate that GLUT1/3-mediated uptake of extracellular glucose is indispensable for neutrophil glycolysis, NADPH generation via the PPP, ROS production and secretion of pro-inflammatory LMs. Notably, loss of GLUT1/3 function abrogates the oxidative burst and COX-dependent PG synthesis, while promoting the production of immunomodulatory metabolites and LMs. These findings position glucose uptake as a central metabolic checkpoint that links pathogen sensing to the inflammatory phenotype of murine and human neutrophils.

Different tissues and cell types express a heterogeneous array of hexose transporters (Mueckler and Thorens, 2013), with neutrophils predominantly expressing the high-affinity glucose transporters GLUT1 and GLUT3 (Li et al., 2022). While the role of GLUT1 in neutrophils has been explored in some studies (Li et al., 2022, Sun et al., 2025, Schuster et al., 2007, Ancey et al., 2021, Britt et al., 2024), the specific contribution of GLUT3 to glucose uptake, glycolytic metabolism and antimicrobial effector function remains poorly understood. Interestingly, stimulation of murine and human neutrophils with *C. albicans* or *S. aureus* markedly increased the expression of GLUT1 at both mRNA and protein levels, whereas GLUT3 expression was unaltered or downregulated. These findings align with previous studies reporting elevated GLUT1 expression and glucose utilization in response to fungal, bacterial and tumor antigens (Li et al., 2022, Ancey et al., 2021, Schuster et al., 2007). However, genetic deletion of GLUT1 alone led to only a modest reduction in glucose uptake and glycolytic flux in neutrophils (Li et al., 2022), suggesting that GLUT3 may play a predominant role in neutrophil glucose metabolism. Supporting this notion, combined deletion of GLUT1 and GLUT3 almost completely abolished glucose uptake and glycolysis upon microbial stimulation. Remarkably, however, despite profound disruption of a key metabolic pathway, loss of both transporters did not impair neutrophil development or alter circulating neutrophil counts. These findings suggest that homeostatic granulopoiesis and neutrophil survival are largely independent of extracellular glucose and may instead rely on alternative metabolic pathways, such as glutaminolysis, lipolysis or mitochondrial respiration (Ettel and Weichhart, 2024, Thind et al., 2024). Furthermore, both murine and human neutrophils harbor substantial cytosolic glycogen stores that can sustain glycolytic metabolism even under conditions of limited extracellular glucose availability (Sadiku et al., 2017, Sadiku et al., 2021, Britt et al., 2024). This observation raises the possibility that endogenous glycogen may support neutrophil development in the absence of GLUT1/3-mediated glucose uptake under steady-state conditions. However, whether extracellular glucose is required to replenish these glycogen stores and sustain emergency and extramedullary granulopoiesis during inflammation remains an open question and warrants further investigation.

In contrast to homeostatic conditions, acute stimulation of neutrophils induces a substantial increase in extracellular glucose utilization to fuel their antimicrobial functions (Britt et al., 2024, Lika et al., 2025, Li et al., 2022). Here, we show that glucose uptake in pathogen-stimulated neutrophils is almost exclusively mediated by GLUT1 and GLUT3, with minimal contribution or compensatory activity from other hexose transporters. Once inside the cytoplasm, glucose is rapidly phosphorylated by hexokinases to glucose-6-phosphate (G6P), which prevents it diffusion out of the cell. In activated neutrophils, G6P is primarily converted into lactate – even under aerobic conditions – rather than being oxidized in the mitochondria via the TCA cycle (Jeon et al., 2020, Lika et al., 2025). In addition, G6P also fuels the PPP, particularly its oxidative branch, to generate large amounts of NADPH. This reducing equivalent is essential for various biosynthetic processes and – especially in neutrophils – for the activation of the NADPH oxidase complex. Within this multi-enzyme complex, NADPH donates electrons for the rapid reduction of molecular oxygen to superoxide and other ROS, driving the respiratory burst that enables neutrophils to kill phagocytosed pathogens though oxidative damage (Britt et al., 2022). The importance of glucose-dependent NADPH production is underscored by patients with G6P dehydrogenase deficiency. In these individuals, impaired NADPH generation compromises the oxidative burst and neutrophil function, leading to recurrent bacterial and fungal infections (Siler et al., 2017). Similarly, ablation of GLUT1 and GLUT3 significantly impaired the oxidative branch of the PPP and NADPH-dependent ROS production upon stimulation with *C. albicans* and *S. aureus*, highlighting the dependence of the respiratory burst on extracellular glucose in murine and human neutrophils. Intriguingly, *de novo* synthesis of purine nucleotides and nucleosides was not completely abolished in GLUT1/3-deficient neutrophils, indicating an imbalance between the oxidative and non-oxidative branches of the PPP when extracellular glucose uptake is restricted. Moreover, mitochondrial respiration and the activity of the TCA cycle remained largely intact, suggesting that mitochondrial metabolism may partially compensate for the bioenergetic deficit in the absence of GLUT1/3-mediated glucose uptake. Notably, GLUT1/3-deficient neutrophils exhibited elevated levels of the TCA cycle metabolite itaconate, indicating a shift toward an immunomodulatory metabolic program when extracellular glucose utilization is impaired.

Beyond central carbon metabolism, we also examined the impact of GLUT1/3-mediated glucose uptake on LM production following microbial stimulation. Glycolytic metabolism may support the biosynthesis of PUFAs through glucose-derived acetyl-CoA. This acetyl-CoA can be generated from mitochondrial citrate exported to the cytosol and subsequently converted to acetyl-CoA by ATP citrate lyase (Hochrein et al., 2022, Guertin and Wellen, 2023). However, although acetyl-CoA is essential for glucose-driven lipogenesis of fatty acids, cholesterol and isoprenoids, omega-6 (AA) and omega-3 PUFAs (EPA and DHA) cannot be synthesized *de novo* in humans due to the lack of specific desaturase enzymes, namely delta-12 and delta-15 desaturases, which introduce double bonds at precise positions within the fatty acid chain (D’Angelo et al., 2020). Instead, they are derived from essential dietary lipid precursors — linoleic acid and alpha-linolenic acid, respectively — through a series of desaturation and elongation steps (Mika et al., 2020). Supporting this notion, we observed no incorporation of glucose-derived ^13^C-labels into lipids or their precursors in zymosan-stimulated neutrophils during our metabolic tracing experiments. Nonetheless, genetic or pharmacological suppression of GLUT1/3 significantly altered the LM profile of murine and human neutrophils, suggesting that these effects are mediated through indirect regulatory pathways. Several steps in PUFA and LM metabolism rely on ATP and NADPH – key metabolites generated through glycolysis and the PPP (Smyrniotis et al., 2014). Moreover, altered bioenergetics or disrupted signaling pathways in the absence of GLUT1/3-mediated glucose consumption, such as impaired mTOR signaling (Weichhart et al., 2015), disrupted redox homeostasis (Braunstein et al., 2025) and perturbations in post-translational protein modifications (Thimmappa et al., 2024), may further contribute to the altered LM production in GLUT1/3-deficient neutrophils. While stimulation with *S. aureus* induced only a modest LM production by murine and human neutrophils, fungal antigens elicited a robust release of diverse LM species. Notably, the production of PGE_2_, PGF_2α_, TXB_2_, as well as several monohydroxylated eicosanoid and docosahexaenoic-derived lipid species was significantly reduced in the absence of GLUT1 and GLUT3. By contrast, levels of LTB_4_ and dihydroxylated eicosanoid acids, such as 5S,6R-diHETE, were significantly elevated in GLUT1/3-deficient neutrophils. These shifts in LM production were also observed in human neutrophils treated with a pharmacological GLUT1/3 inhibitor, demonstrating that glucose metabolism regulates the balance between COX and LOX pathways in both murine and human neutrophils. The increase in LTB_4_ observed in neutrophils with impaired GLUT1/3 activity is particularly noteworthy, as patients with chronic granulomatous disease (CGD) – caused by defects in the NADPH oxidase complex – also exhibit elevated LTB_4_ production in response to fungal antigens (Song et al., 2020). This parallel suggests that altered NADPH-mediated ROS production and cellular redox homeostasis may indirectly influence 5-LOX and LTA_4_ hydrolase activity.

From a translational perspective, our findings are significant because GLUT1 and GLUT3 represent promising therapeutic targets for inflammatory and autoimmune diseases (Chen et al., 2023, Reckzeh and Waldmann, 2020). While suppression of GLUT1 and/or GLUT3 could effectively ameliorate immunopathology in T cell-mediated diseases (Hochrein et al., 2022, Macintyre et al., 2014), the critical role of GLUT1/3-mediated glycolysis in neutrophil-mediated antimicrobial immunity raises concerns about potential adverse effects, such as increased susceptibility to infections. Therefore, the therapeutic benefits of novel GLUT inhibitors must be carefully balanced against the essential roles of GLUT1 and GLUT3 in innate immunity and host defense.

## ACKNOWLEDGEMENTS

We thank Miriam Eckstein for excellent technical assistance, Prof. Dr. Tim Lämmermann (Max Planck Institute of Immunobiology and Epigenetics, Freiburg, Germany) for providing the *Mrp8*^Cre^ mice and Prof. Dr. Herbert Waldmann (Max Planck Institute of Molecular Physiology, Dortmund, Germany) for his sustained support with GLUT1/3 inhibitors. Design, generation and optimization of specific GLUT1 and GLUT3 inhibitors, such as LDC9620 was financed by the Max-Planck Gesellschaft e.V. as part of a translational and collaborative program agreement between the Max-Planck Institute of Molecular Physiology under the guidance and supervision of Prof. Dr. Herbert Waldmann and the Lead Discovery Center GmbH (Dortmund, Germany). The LDC thanks Dr. Lidia Urbina for her critical review of this manuscript. This work was funded by the Deutsche Forschungsgemeinschaft (DFG) SFB-TR 124/3 (“FungiNet”) – project number: 210879364; SFB 1526/1 (“PANTAU”), project number: 454193335; SFB-TR 338/1 (“LETSimmun”) – project number: 452881907, SFB 1583/1 (“DECIDE”) – project number: 49262049; SFB 1127/3 (“ChemBioSys”) – project number: 239748522 and individual project grants VA882/2-1 and VA882/3-2. Further support was provided by the SFB 1525/1 (“Cardio-Immune Interfaces”) – project number: 453989101 and the Mildred Scheel Early Career Center (MSNZ) of the German Cancer Aid. We thank the Free State of Thuringia (Thüringer Aufbaubank) and the European Union (Europäischer Fonds für regionale Entwicklung, 2023 FGI 0012) for financial support.

## AUTHOR CONTRIBUTION

Participated in research design: A.F., M.V., P.M.J, W.S., K.G. and J.M., K.O. and O.W. Conducted experiments: A.F., K.G., M.C.P. S.M.H., W.S., K.S., and P.M.J. Performed data analysis: A.F., K.G., W.S., G.Z., B.K. and P.M.J. Compound design, synthesis and profiling: G.Z. and B.K. Wrote the manuscript: M.V. and A.F.

## METHODS

### Animals

All mice were bred and maintained under specific pathogen free conditions at the Institute of Systems Immunology or the Center for Experimental Medicine (ZEMM) at the Julius-Maximilians-University of Würzburg. Mice were maintained on a 12/12 h light/dark cycle between 20-24°C in individually ventilated cages. Mice had access to standard chow (Ssniff; cat# V1534) and autoclaved water ad libitum and health status of the animals was inspected daily by the responsible animal caretakers. Hygiene status of the sentinel mice was monitored quarterly according to the FELASA guidelines. Both male and female mice between 8 and 28 weeks of age at the time of the experiment were used. C57BL/6J *wild type* (strain 000664), and *Slc2a1*^fl/fl^ (strain 031871) mice were originally purchased from the Jackson Laboratories (JAX) and maintained at our institution. *Slc2a3*^fl/fl^ mice were generated with the knockout mouse project (KOMP) at UC Davis and have been described before (Fidler et al., 2017). *Mrp8*^Cre^ mice (Passegué et al., 2004) were kindly provided by Tim Lämmermann (Max Planck Institute of Immunobiology and Epigenetics, Freiburg, Germany). All animals used in this study were on a C57BL/6 genetic background.

### Fungal and bacterial cultures

*Candida albicans* strain SC5314 was grown overnight (o/n) in yeast extract-peptone-dextrose (YPD) broth at 30°C with constant shaking. For log-phase cultures, o/n cultures were diluted into fresh medium and grown at 30°C until optical density at 600 nm (OD_600_) was reached. Hyphal growth was induced by culturing *C. albicans* o/n in YPD medium supplemented with 10% FCS at 37°C with constant shaking. MOIs were determined from growth curves generated by measuring the OD_600_ and by plating serial dilutions of o/n cultures. *Staphylococcus aureus* strain HG001 was grown in an o/n culture in lysogeny broth (LB) medium at 37°C with constant shaking. For log-phase culture, overnight culture was diluted into fresh LB medium and grown at 37°C and harvested at desired OD_600_. For experiments utilizing inactivated pathogens, *C. albicans* was incubated with 0.1 mg/ml nystatin (Carl Roth) for 60 min and *S. aureus* with 100 units/ml penicillin/streptomycin (Gibco). Inactivation of pathogens was confirmed by plating the treated pathogens on appropriate culture plates.

### Murine and human neutrophil isolation

Bone marrow neutrophils isolated from mouse femurs and tibiae were isolated using the Neutrophil Isolation Kit (Miltenyi Biotec) following the manufacturer instructions. Purity of isolated bone marrow neutrophils was verified by flow cytometry (live Ly6G^+^ CD11b^+^). Human neutrophils were isolated from fresh whole blood samples of healthy donors using MACSxpress Whole Blood Neutrophil Isolation Kit (Miltenyi Biotec) following manufacturer instructions. Purity of human peripheral blood neutrophils was determined by flow cytometry (live CD16^+^ CD11b^+^).

### Stimulation of neutrophils with live and inactivated pathogens

Freshly isolated murine and human neutrophils were rested for 30 min after isolation before every experiment. Neutrophils were cultured and stimulated in modified RPMI 1640 medium containing 100 mg/dl glucose supplemented with 10% fetal calf serum (Sigma) and 0.1% 2-mercaptoethanol (Gibco) at 37°C. Stimulation was performed with live *C. albicans* (strain SC5314) yeast/hyphae or *S. aureus* (strain HG001) from fresh cultures in multiplicity of infection (MOI) of 0.4, 1 or 5, as indicated in the figure legends. Therefore, murine and human neutrophils were seeded in 96-well round bottom plates with a concentration of 5 x10^5^/well and incubated for 3 h (human) or 6 h (mouse) with live pathogens. Human neutrophils were incubated with or w/o 40 mM 2-DG (Sigma) or 4 µM GLUT1/3-Inhibitor LDC9620 (Lead discovery Center GmbH). Assays with inactivated pathogens (MOI = 5) were performed in modified RPMI 1640 containing additionally 100 units/ml penicillin (Gibco) and 100 units/ml streptomycin (Gibco). In metabolomic tracing, murine neutrophils were stimulated with 100 µg/ml *Saccharomyces cerevisiae* zymosan A (Sigma Aldrich) in glucose-free RPMI 1640 containing for 100 mg/dl [U-^13^C_6_]-glucose (Sigma) supplemented with 10% fetal calf serum (Sigma) and 0.1% 2-mercaptoethanol (Gibco) for 12 h.

### Flow cytometric analyses

Flow cytometry analysis was conducted as previously described (Wu et al., 2023). In brief, cells were stained with fluorophore-conjugated antibodies (**Table S3**) and Fixable Viability Dye eFluor 780 (eBioscience) for 15 min at RT in PBS. After washing with PBS containing 0.5% BSA, cells were fixed using the IC Fixation buffer (eBioscience) for 30 min. For intracellular staining, cells were washed with Permeabilization Buffer (eBioscience) according to manufacturer instructions and subsequently stained o/n with unconjugated antibodies at 4°C in permeabilization buffer (eBioscience). Detection of primary antibodies was carried out with fluorophore-conjugated secondary antibodies for 40 min at RT. All samples were acquired on a BD Celesta flow cytometer (BD Biosciences) or a spectral Aurora Flow Cytometer (Cytek) and analyzed using FlowJo software (BD Biosciences).

### Quantitative real-time PCR and PrimeFlow RNA analysis

Total RNA from resting and stimulated murine neutrophils was extracted using Roti-Prep Mini Kit (Roth) and mRNA concentration quantified via Nanodrop analysis (Thermo Fisher Scientific). The cDNA synthesis was performed using iScript cDNA synthesis kit (Bio-Rad) according to manufacturer’s instructions. The transcript levels were measured using the SYBR Green qPCR MasterMix (Bio-Rad) and primers for *Slc2a1* (F: GAGACCAAAGCGTGGTGAGT, R: GCAGTTCGGCTATAACACTGG) and *Slc2a3* (F: ATCGTGGCATAGATCGGTTC, R: TCTCAGCAGCTCTCTGGGAT). The ribosomal protein 18S (F: CGGCGACGACCCATTCGAAC, R: GAATCGAACCCTGATTCCCCGT) was used as a reference gene for normalizing the samples via the 2^-ΔCT^ method. Intracellular human *SLC2A1* and *SLC2A3* mRNA expression was detected using the PrimeFlow RNA Assay Kit (Thermo Fisher Scientific), according to the manufacturer’s instructions. Following cell surface staining of protein antigens with fluorochrome-conjugated antibodies, probe-hybridization was performed at 40°C for 1 h. Commercially available, pre-designed RNA probes specific for human *SLC2A1* and *SLC2A3* (Thermo Fisher Scientific) were hybridized with PreAmplifier and Amplifier probes for 90 min at 40°C followed by incubation with fluorochrome-conjugated Label-probes conjugated with Alexa Fluor 488 (Type 1) for 60 min. Detection of *SLC2A1* and *SLC2A3* mRNA at single-cell level was carried out using flow cytometry.

### Radiolabeled glucose uptake assay

To measure glucose uptake of neutrophils, 2.5x10^5^ murine and human neutrophils were stimulated in a 96-well plate with inactivated *C. albicans* (0.1 mg/ml nystatin) or *S. aureus* (100 units/ml penicillin/streptomycin) at a MOI = 5 for 3h (human) or 6h (mouse). Stimulation was performed in glucose-free RPMI 1640 medium (Thermo Fisher Scientific) supplemented with 1 μCi/ml [^3^H]-2-deoxy-D-glucose ([^3^H]-2-DG) (Perkin Elmer), 10% fetal calf serum (Sigma), 100 units/ml penicillin/streptomycin, 0.1% 2-mercaptoethanol (Gibco) at 37°C. Cells were washed twice with glucose-free RPMI medium and resuspended in 140 μl water to lyse the cells. 40 μl of this lysate was added to 160 μl scintillation cocktail and [^3^H] counts per minute (cpm) were measured in triplicates using a MicroBeta2 microplate scintillation counter (Perkin Elmer).

### Seahorse extracellular flux analysis

Lactate secretion and mitochondrial respiration in neutrophils were assessed via glycolytic proton efflux rate (glycoPER) and oxygen consumption rate (OCR), respectively, using an oxygen-controlled XFe96 extracellular flux analyzer (Seahorse Bioscience) as previously described (F. Gubert et al., 2025). 3x10^5^ neutrophils per well were seeded on Cell-Tak (Corning) coated XFe96 cell culture microplate (Agilent) in standard Seahorse XF RPMI medium (Agilent), supplemented with 2 mM L-glutamine (Thermo Fisher Scientific), 1 mM sodium pyruvate (Sigma), 10 mM D-glucose (Sigma), 100 units/ml penicillin and 100 units/ml streptomycin (both Gibco). Murine neutrophils were stimulated with inactivated *C. albicans* (0.1 mg/ml nystatin) or *S. aureus* (100 units/ml penicillin/streptomycin) for 6 h at 37°C in a non-CO2-incubator. Human neutrophils were incubated with or w/o 40 mM 2-DG (Sigma) or 4 µM GLUT1/3-Inhibitor LDC9620 (Lead discovery Center GmbH) for 3 h at 37°C. LDC9620 is a representative tool compound of a larger series of GLUT1- and GLUT3-selective inhibitors, which have been designed and synthesized at the LDC. The chemical structure of the compounds will be published elsewhere. Research amounts of LDC9620 can be provided by LDC upon request. After stimulation, neutrophils were supplemented with fresh Seahorse assay medium and metabolic stress tests were performed following the manufacturer’s guidelines. For assessing basal glycolytic rates, the extracellular acidification rate (ECAR) was measured. By adding 5 µM Rotenone (Biomol) and 5 µM Antimycin A (Sigma) mitochondrial complexes I and III were inhibited to measure compensatory glycolysis. To block glycolysis completely 100 mM 2-DG (Sigma) were added. To calculate the glycolytic proton efflux rate (PER) the buffer factor of the medium was included (F. Gubert et al., 2025). Mitochondrial stress tests were performed to evaluate basal and maximal mitochondrial respiration. Basal oxygen consumption was measured, before the addition of 2 µM oligomycin (Cayman Chemicals) to calculate the ATP production rate. 1 µM FCCP (Cayman Chemicals), was added to induce maximal respiration. To calculate the spare respiratory capacity (SRC), 5 µM rotenone (Biomol) and 0.5 µM antimycin A (Sigma) were added.

### ROS production assay

To measure the production of intra- and extracellular ROS, freshly isolated murine and human neutrophils were resuspended in phenol red-free RPMI 1640 medium supplemented with 1% fetal calf serum (Sigma), 25 mM HEPES, 2g/l NaHCO_3_, 2 mM glutamine and 40 µg/ml luminol (Biomol). A total of 2x10^5^ neutrophils per well were seeded into white 96-well plates (Corning) and stimulated with *C. albicans* or *S. aureus* (both at MOI = 1) immediately prior to measurement. Luminescence was recorded over 6 h at 37°C using a FlexStation III plate reader (Molecular devices) and displayed as relative light units (RLU).

### Metabolomic tracing using LC-MS

To analyze total and ^13^C-labeled polar metabolites, murine bone marrow neutrophils were incubated with 100 mg/dl [U-^13^C_6_]-glucose (Sigma) in glucose-free RPMI 1640 medium supplemented with 10% fetal calf serum (Sigma) and 0.1% 2-mercaptoethanol (Gibco). Neutrophils were stimulated with 100 µg/ml zymosan A (Sigma Aldrich) for 12 h at 37°C. Following stimulation, cells were harvested, and the resulting cell pellets were washed with ice-cold 0.9% NaCl and then snap-frozen in liquid nitrogen. Subsequent polar metabolite extraction, sample acquisition and data analysis was performed as described before (Wu et al., 2023). In brief, cells were resuspended in 360 µl 10 mM HCl/MeOH (17/19, v/v) containing external water-soluble standard compounds (0.02 µM lamivudine and 2 µM each of D_2_-glucose, D_4_-succinate, D_5_-glycine and ^15^N-glutamate) and treated with ultrasound (Branson). After vigorous mixing, another 100 µl CHCl_3_ were added, and the resulting phases separated after mixing and centrifugation. The upper phase was evaporated under a stream of nitrogen for 15 min and taken to dryness in a centrifugal evaporator. Liquid chromatography and mass spectrometry (LC-MS) analyses were performed on a Dionex Ultimate 3000 UHPLC system connected to a Q Exactive mass spectrometer (QE-MS) equipped with a HESI probe (both Thermo Fischer Scientific). Chromatographic separation was achieved by applying 3 µl sample on a XBridge Premier BEH Amide (100 x 2.1 mm, 2.5 μm) (Waters), protected by a Supelco ColumnSaver particle filter (Merck) and a gradient of mobile phase A (5 mM NH_4_OAc in CH_3_CN/H_2_O: 40/60, v/v) and mobile phase B (5 mM NH_4_OAc in CH_3_CN/H_2_O: 95/5, v/v) maintaining a flow rate of 200 µl/min. The LC gradient program was 100% mobile phase B for 2 min, followed by a linear decrease to 20% B within 23 min, maintaining 20% B for 21 min and returning to 100% B in 2 min, followed by 7 min 100% B for column equilibration before each injection. The eluent was directed to the QE-MS from 2.7 min to 46 min after sample application. Mass detection was conducted in alternating pos/neg full scan mode (at 70k resolution, scan range m/z 69 - 1000, AGC target 1E6 and 200 ms max. injection time). HESI parameters: Sheath gas: 20, aux gas: 1, spray voltage: 3.0 kV, capillary temp.: 300 °C, S-lens RF level: 50.0, aux gas heater temperature: 120 °C.

### Lipid mediator metabololipidomics using UPLC-MS/MS

Freshly isolated murine and human neutrophils were cultured in modified phenol red-free RPMI 1640 medium supplemented with 10% fetal calf serum (Sigma) and 0.1% 2-mercaptoethanol (Gibco) at a concentration of 5 x10^5^/well in 96-well round bottom plates. Cells were stimulated with live *C. albicans* (hyphae) and *S. aureus* (both at MOI = 1) for 3 h (human) or 6 h (mouse) at 37° C. The supernatants of the cells were collected and solid phase extraction (SPE) and sample preparation for UPLC-MS/MS analysis was conducted by adapting published criteria (Werz et al., 2018). Briefly, samples were kept at -20°C for at least 60 min to allow protein precipitation. After centrifugation (1200 g, 4°C, 10 min), 9 ml acidified H_2_O was added to the supernatants (pH = 3.5) and samples were subjected to SPE. Solid phase cartridges (Sep-Pak Vac 6cc 500 mg / 6 ml C18; Waters) were equilibrated with 6 ml methanol and 2 ml H_2_O. Then, samples were loaded onto columns and washed with 6 ml H_2_O and additional 6 ml n-hexane. LM were eluted with 6 ml methyl formate, the eluates were brought to dryness using an evaporation system (TurboVap LV, Biotage), and LM were resuspended in 100 µl methanol/water (50:50, v/v) for UPLC-MS/MS analysis. LM profile was analyzed with an Acquity UPLC system (Waters) and a QTRAP 5500 Mass Spectrometer (ABSciex) equipped with a Turbo V Source and electrospray ionization. LM were separated using an ACQUITY UPLC BEH C18 column (1.7 µm, 2.1x100 mm; Waters) at 50°C with a flow rate of 0.3 ml/min and a mobile phase consisting of methanol/water/acetic acid of 42/58/0.01 (v/v/v) that was ramped to 86/14/0.01 (v/v/v) over 12.5 min and then to 98/2/0.01 (v/v/v) for 3 min (Werner et al., 2019). The QTrap 5500 was operated in negative ionization mode using scheduled multiple reaction monitoring coupled with information-dependent acquisition. The scheduled multiple reaction monitoring window was 60 sec, and the curtain gas pressure was set to 35 psi. The retention time and at least six diagnostic ions for each LM were confirmed by means of an external standard (Cayman Chemical). Quantification was achieved by calibration curves for each LM. Linear calibration curves were obtained for each LM and gave r2 values of 0.998 or higher. Additionally, the limit of detection for each targeted LM was determined (Werner et al., 2019). The identity of low abundance analytes was confirmed by fragmentation pattern matching by re-analysis using a QTrap 7500 mass spectrometer (Sciex) controlled by SCIEX-OS and comparing the enhanced product ion scans of the biological sample with that of authentic standards.

### Quantification and statistical analysis

The results are shown as mean ± standard error of the means (SEM). To determine the statistical significance of the differences between the experimental groups unpaired or paired Student’s t tests were performed using the Prism 10 software (GraphPad). Sample sizes were based on experience and experimental complexity. Differences reached significance with p values < 0.05 (noted in figures as *), p < 0.01 (**), p < 0.001 (***) and ****p < 0.0001. The figure legends contain the number of independent samples or mice per group and the statistical analyses that were used.

