## Supplementary material for "Glucose Metabolism Controls Oxidative Burst and Lipid Mediator Production in Neutrophils upon Microbial Challenge": SUPPLEMETARY INFORMATION

#### Supplementary Figures S1–S4

#### Supplementary Tables S1 – S3

### Supplementary Figures S1

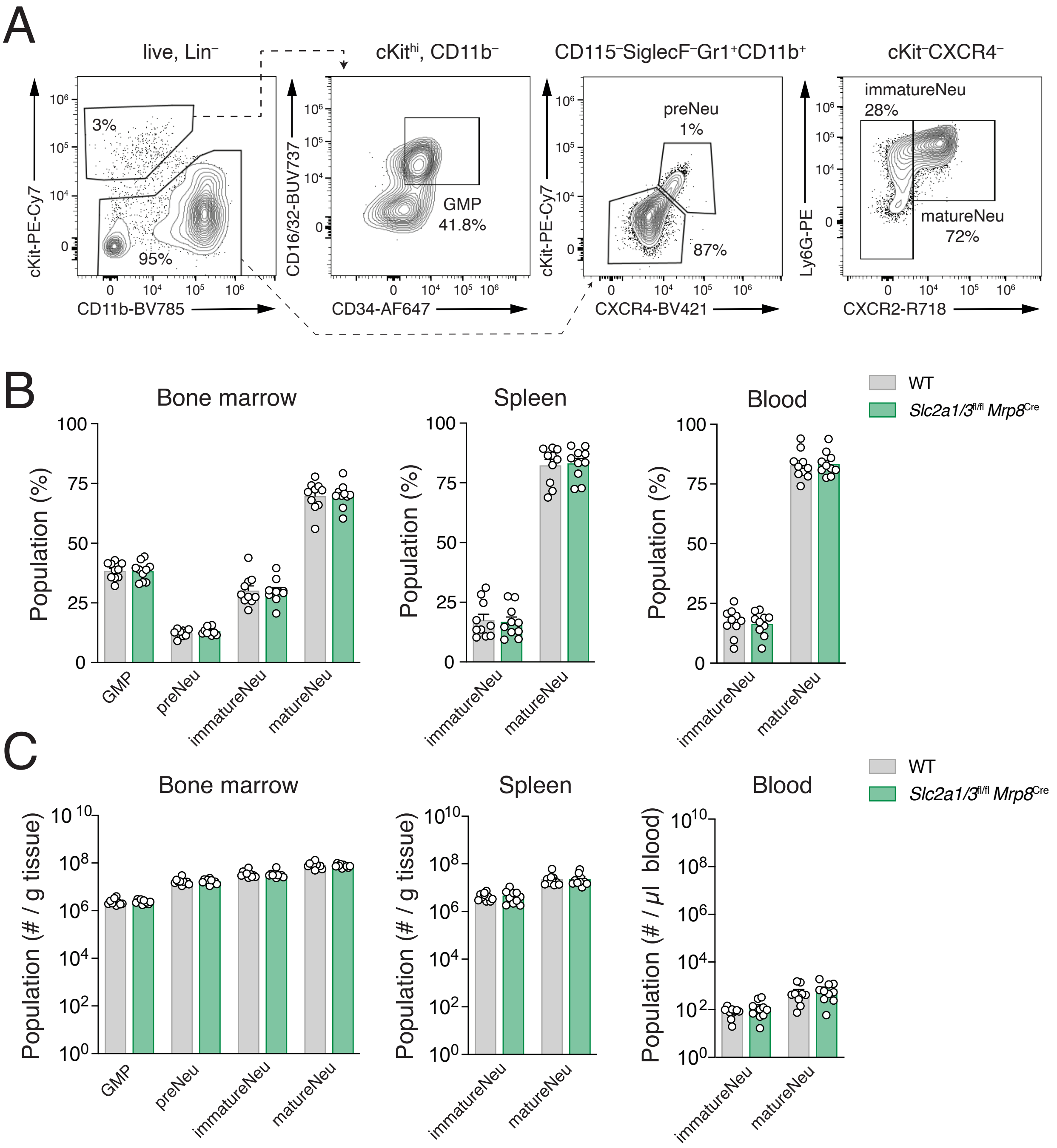

**Figure S1. GLUT1 and GLUT3 are dispensable for neutrophil development and activation. (A)** Gating strategy to analyse neutrophil development in bone marrow (BM), spleen and blood. Lineage negative (CD3ε⁻, NK1.1⁻, B220⁻, Sca-1⁻, CD90.2⁻) viable singlet cells were gated according to scheme to define granulocyte-monocyte progenitors (GMPs), committed neutrophil precursors (preNeu), immature neutrophils (immatureNeu) and mature neutrophils (matureNeu). **(B)** Quantification of neutrophil populations in BM, spleen and blood in WT and GLUT1/3-deficient (*Slc2a1*$^{fl/fl}$ *Slc2a3*$^{fl/fl}$ *Mrp8*$^{Cre}$) mice as described in (A); means ± SEM of 10 mice. **(C)** Absolute cell counts of neutrophil populations in BM, spleen and blood in WT and GLUT1/3-deficient mice; cell numbers were normalized to organ weight or blood volume; means ± SEM of 10 mice.

### Supplementary Figures S2

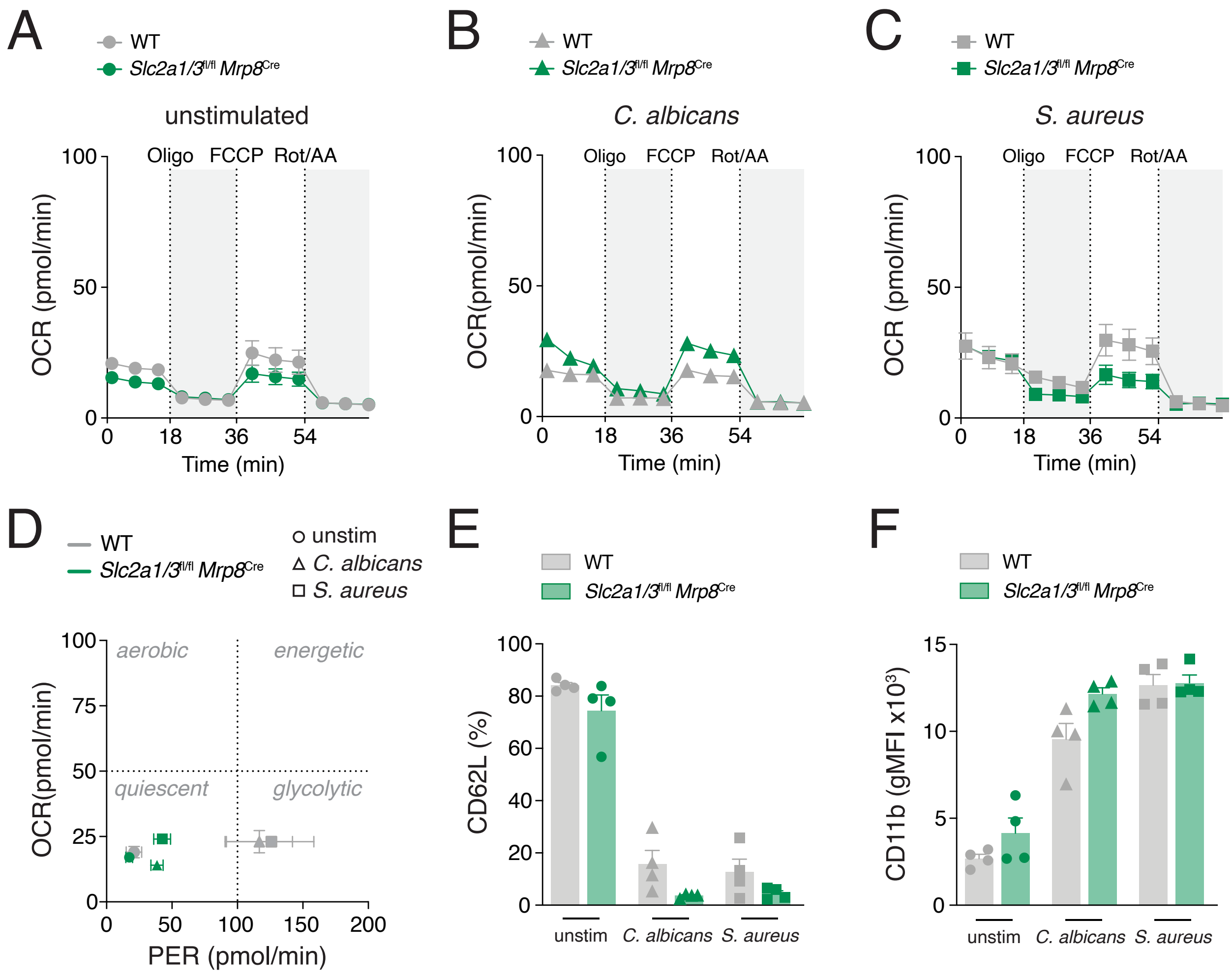

**Figure S2. Mitochondrial respiration in neutrophils following *C. albicans* and *S. aureus* stimulation.** (**A**) Analysis of oxygen consumption rate (OCR) in WT and GLUT1/3-deficient (*Slc2a1*$^{fl/fl}$ *Slc2a3*$^{fl/fl}$ *Mrp8*$^{Cre}$) neutrophils left unstimulated **(A)** or stimulated with inactivated *C. albicans* (hyphae) **(B)** or *S. aureus* **(C)** (both MOI = 5) for 6 h using a Seahorse extracellular flux analyser; means ± SEM of 4 biological replicates per group. **(D)** Energy phenotype profiling of WT and GLUT1/3-deficient neutrophils left unstimulated or stimulated for 6 h with inactivated *C. albicans* (hyphae) or *S. aureus* as described in (A-C); Oxygen consumption rate (OCR) vs. glycolytic proton efflux rate (PER); mean ± SEM of 4 biological replicates per group. **(E** and **F)** Flow cytometric analysis of CD62L (**E**) and CD11b (**F**) expression on WT and GLUT1/3-deficient neutrophils stimulated with live *C. albicans* (hyphae) or *S. aureus* (both MOI = 1) for 6 h; means ± SEM of 4 biological replicates per group.

### Supplementary Figures S3

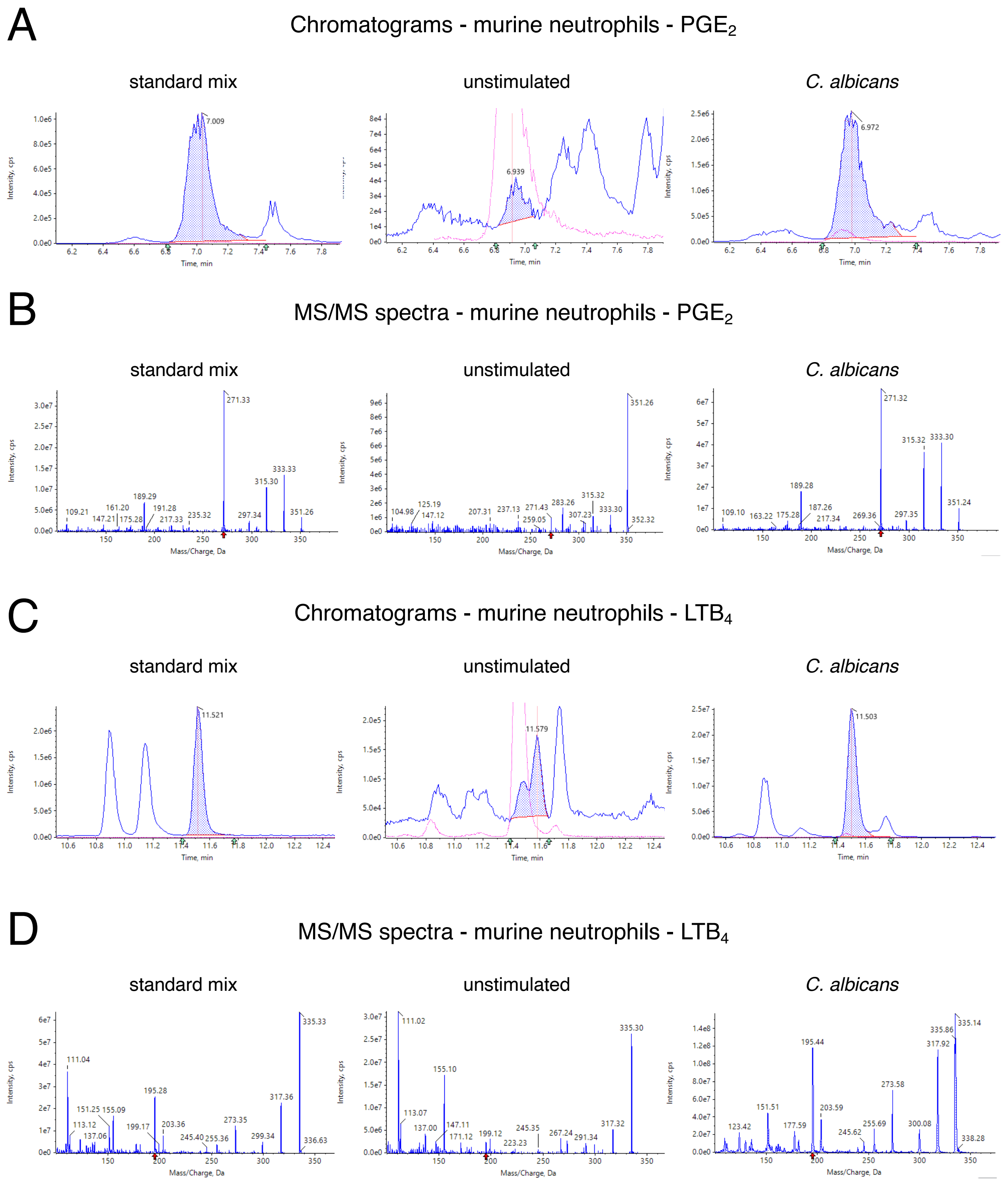

**Figure S3. Representative chromatograms and mass spectra of LMs secreted by murine neutrophils.** (**A-D**) Representative chromatograms and mass spectra of $PGE_2$ (A and B) and $LTB_4$ (C and D) from standard mix and cell culture supernatants of murine bone marrow WT neutrophils treated with or without *C. albicans* (hyphae) for 6 h (MOI = 1) analyzed by ultra performance liquid chromatography – tandem and mass spectrometry (UPLC-MS/MS); n.d. = not detected.

### Supplementary Figures S4

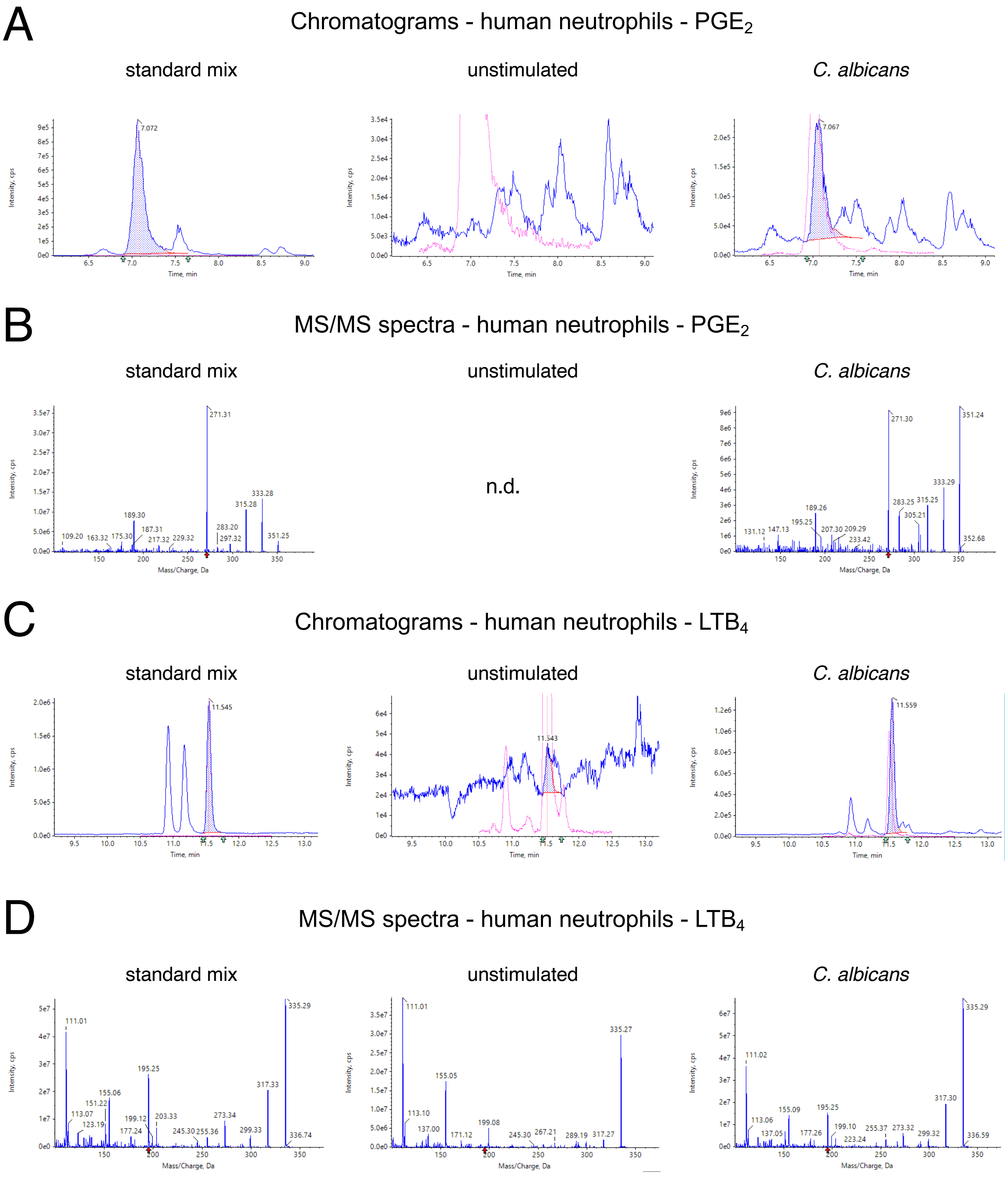

**Figure S4. Representative chromatograms and mass spectra of LMs secreted by human neutrophils.** (**A-D**) Representative chromatograms and mass spectra of $PGE_2$ (A and B) and $LTB_4$ (C and D) from standard mix and cell culture supernatants of human peripheral bold neutrophils treated with or without *C. albicans* (hyphae) for 6 h (MOI = 1) analyzed by ultra performance liquid chromatography – tandem and mass spectrometry (UPLC-MS/MS); n.d. = not detected.

### Supplementary Table S1

| | | unstim | | | *C. albicans* | | | | *S. aureus* | | | |
|---|---|---|---|---|---|---|---|---|---|---|---|---|
| | | WT | *Slc2a1/3*[fl/fl] *Mrp8*[cre] | | WT | | *Slc2a1/3*[fl/fl] *Mrp8*[cre] | | WT | | *Slc2a1/3*[fl/fl] *Mrp8*[cre] | |
| | pg/2.5x10[6] cells | | | fold | | fold | | fold | | fold | | fold |
| COX | $PGD_1$ | <10 | <10 | 1,00 | 131,34 ± 34,42 | 13,13 | 129,89 ± 21,11 | 12,99 | 85,27 ± 10,02 | 8,53 | 96,51 ± 13,34 | 9,65 |
| | $PGD_2$ | 243,52 ± 61,93 | 249,56 15,84 | 1,02 | 1694,68 ± 196,48 | 6,96 | 1561,91 ± 100,30 | 6,41 | 667,46 ± 39,85 | 2,74 | 886,40 ± 158,81 | 3,64 |
| | $PGE_2$ | 32,51 ± 8,18 | 40,89 ± 3,01 | 1,26 | 3974,44 ± 421,52 | 122,26 | 3247,40 ± 121,88 | 99,90 | 1421,86 ± 207,93 | 43,74 | 1623,20 ± 180,79 | 49,93 |
| | $PGF_{2a}$ | <10 | <10 | 1,00 | 1463,93 ± 182,44 | 146,39 | 1089,68 ± 94,09 | 108,97 | 697,90 ± 107,13 | 69,79 | 745,24 ± 55,73 | 74,52 |
| | $TXB_2$ | 561,00 ± 62,02 | 582,04 ± 32,20 | 1,04 | 6483,67 ± 588,45 | 11,56 | 4468,59 ± 428,25 | 7,97 | 2265,02 ± 431,96 | 4,04 | 2107,62 ± 200,68 | 3,76 |
| 5-LOX | 5-HETE | 1207,19 ± 56,94 | 1290,09 ± 95,15 | 1,07 | 9552,42 ± 879,38 | 7,91 | 10132,28 ± 1488,74 | 8,39 | 1788,49 ± 133,58 | 1,48 | 2356,71 ± 230,34 | 1,95 |
| | 5-HEPE | 233,18 ± 10,45 | 241,53 ± 1,14 | 1,04 | 363,27 ± 11,81 | 1,56 | 348,86 ± 30,17 | 1,50 | 221,62 ± 15,81 | 0,95 | 239,43 ± 18,81 | 1,03 |
| | 7-HDHA | 330,87 ± 22,26 | 323,04 ± 13,77 | 0,98 | 531,05 ± 46,98 | 1,61 | 528,08 ± 56,51 | 1,60 | 336,75 ± 9,04 | 1,02 | 338,76 ± 13,40 | 1,02 |
| | t-$LTB_4$ | 50,28 ± 2,87 | 53,88 ± 1,78 | 1,07 | 8175,48 ± 470,43 | 162,61 | 5579,15 ± 783,24 | 110,97 | 219,15 ± 22,94 | 4,36 | 365,84 ± 109,92 | 7,28 |
| | et-$LTB_4$ | 161,28 ± 4,98 | 177,88 ± 10,90 | 1,10 | 3615,09 ± 241,14 | 22,41 | 3527,84 ± 619,99 | 21,87 | 316,91 ± 24,10 | 1,96 | 441,83 131,34 | 2,74 |
| | $LTB_4$ | 107,40 ± 6,36 | 114,96 ± 3,58 | 1,07 | 15580,28 ± 3154,05 | 145,06 | 23390,62 ± 1662,76 | 217,78 | 1348,25 ± 163,17 | 12,55 | 2441,07 ± 578,77 | 22,73 |
| | $LTB_5$ | <10 | <10 | 1,00 | 179,16 ± 32,87 | 17,92 | 288,73 ± 25,94 | 28,87 | 16,33 ± 1,27 | 1,63 | 25,84 ± 5,68 | 2,58 |
| | 5,6-diHETE | 14,98 ± 1,00 | 15,56 ± 1,39 | 1,04 | 943,57 ± 84,86 | 63,00 | 1777,59 ± 287,11 | 118,68 | 134,11 ± 9,45 | 8,95 | 222,29 ± 53,29 | 14,84 |
| monohydroxylated FA | 15-HETE | 547,81 ± 29,64 | 552,90 ± 15,07 | 1,01 | 1083,99 ± 11,32 | 1,98 | 789,62 ± 28,71 | 1,44 | 618,17 ± 40,09 | 1,13 | 635,40 ± 36,10 | 1,16 |
| | 11-HETE | 258,49 ± 16,23 | 251,70 ± 0,85 | 0,97 | 811,65 ± 35,24 | 3,14 | 603,82 ± 52,56 | 2,34 | 370,38 ± 4,69 | 1,43 | 396,95 ± 7,93 | 1,54 |
| | 8-HETE | 150,96 ± 4,23 | 142,05 ± 2,57 | 0,94 | 229,52 ± 19,80 | 1,52 | 215,88 ± 23,50 | 1,43 | 153,86 ± 6,90 | 1,02 | 151,96 ± 4,16 | 1,01 |
| | 5,15-diHETE | 108,23 ± 3,34 | 112,17 ± 4,58 | 1,04 | 328,51 ± 26,80 | 3,04 | 368,26 ± 30,79 | 3,40 | 120,17 ± 5,94 | 1,11 | 140,89 ± 10,81 | 1,30 |
| | 13-HODE | 237,36 ± 22,77 | 232,32 ± 3,55 | 0,98 | 1053,79 ± 138,83 | 4,44 | 1121,49 ± 92,46 | 4,72 | 255,79 ± 24,83 | 1,08 | 246,24 ± 6,73 | 1,04 |
| | 9-HODE | 150,58 ± 38,21 | 164,95 ± 2,78 | 1,10 | 315,31 ± 20,65 | 2,09 | 259,23 ± 15,40 | 1,72 | 182,17 ± 19,92 | 1,21 | 170,25 ± 7,53 | 1,13 |

**Table S1. Amounts of lipid mediators secreted by murine neutrophils.** Concentrations of lipid mediators in the supernatant of WT and GLUT1/3-deficient (*Slc2a1*[fl/fl] *Slc2a3*[fl/fl] *Mrp8*[Cre]) murine neutrophils after stimulation with *C. albicans* or *S. aureus* as shown in Figure 3B; concentrations are given as pg / 2.5x10[6] neutrophils. Means ± SD of 3 biological replicates per group.

### Supplementary Table S2

| | | unstim | unstim + LDC9620 | | *C. albicans* | | *C. albicans* + LDC9620 | | *S. aureus* | | *S. aureus* + LDC9620 | |
|---|---|---|---|---|---|---|---|---|---|---|---|---|
| | pg/2.25x10[6] cells | | | fold | | fold | | fold | | fold | | fold |
| COX | $PGD_2$ | 37,51 ± 1,76 | 52,55 ± 2,67 | 1,40 | 121,48 ± 33,13 | 3,24 | 103,15 ± 12,03 | 2,75 | 36,39 ± 10,04 | 0,97 | 59,99 ± 8,65 | 1,60 |
| | $PGE_2$ | <10 | <10 | 1,00 | 148,45 ± 83,56 | 14,85 | 48,18 ± 65,82 | 4,82 | 21,5 ± 14,54 | 2,15 | 10,98 ± 1,55 | 1,10 |
| | $TXB_2$ | 348,34 ± 13,36 | 323,48 ± 12 | 0,93 | 631,58 ± 125,31 | 1,81 | 365,34 ± 33,00 | 1,05 | 377,24 ± 78,98 | 1,08 | 358,46 ± 27,49 | 1,03 |
| 5-LOX | 5-HETE | 165,84 ± 12,25 | 209,38 ± 27,12 | 1,26 | 545,29 ± 291,7 | 3,29 | 732,34 ± 383,39 | 4,42 | 175,62 ± 97,62 | 1,06 | 191,14 ± 10,14 | 1,15 |
| | 5-HEPE | 47,85 ± 2,90 | 55,74 ± 5,90 | 1,17 | 89,6 ± 18,48 | 1,87 | 93,56 ± 14,59 | 1,96 | 47,19 ± 7,25 | 0,99 | 55,06 ± 1,75 | 1,15 |
| | 7-HDHA | 48,37 ± 2,41 | 51,61 ± 3,04 | 1,07 | 96,49 ± 14,35 | 1,99 | 82,09 ± 8,86 | 1,7 | 53,57 ± 7,64 | 1,11 | 60,43 ± 3,27 | 1,25 |
| | t-$LTB_4$ | <10 | <10 | 1,00 | 85,29 ± 75,19 | 8,53 | 118,16 ± 84,49 | 11,82 | 18,72 ± 15,06 | 1,87 | 10,96 ± 2,59 | 1,10 |
| | et-$LTB_4$ | <10 | <10 | 1,00 | 114,24 ± 64,35 | 11,42 | 197,6 ± 102,96 | 19,76 | 27,2 ± 33,03 | 2,72 | 13,62 ± 16,46 | 1,36 |
| | $LTB_4$ | <10 | <10 | 1,00 | 301,94 ± 316,03 | 30,19 | 648,24 ± 505,61 | 64,82 | 118,4 ± 199,74 | 11,84 | 50,16 ± 48,57 | 5,02 |
| | 5,6-diHETE | <10 | <10 | 1,00 | 17,76 ± 10,1 | 1,78 | 38,23 ± 24,79 | 3,82 | 11,43 ± 4,25 | 1,14 | <10 | 1,00 |
| monohydroxylated FA | 15-HETE | 84,54 ± 5,40 | 96,1 ± 7,13 | 1,14 | 199,36 ± 51,64 | 2,36 | 187,33 ± 23,13 | 2,22 | 105,7 ± 18,14 | 1,25 | 140,11 ± 3,35 | 1,66 |
| | 11-HETE | 54,45 ± 2,48 | 54,82 ± 4,09 | 1,01 | 123,17 ± 31,81 | 2,26 | 94,24 ± 11,8 | 1,73 | 64,45 ± 11,24 | 1,18 | 70,62 ± 1,65 | 1,30 |
| | 8-HETE | 32,07 ± 0,90 | 32,52 ± 2,21 | 1,01 | 84,28 ± 23,31 | 2,63 | 55,65 ± 9,19 | 1,74 | 38,83 ± 6,17 | 1,21 | 42,71 ± 1,21 | 1,33 |
| | 18-HEPE | 30,48 ± 1,36 | 31,26 ± 2,08 | 1,03 | 62,1 ± 8,98 | 2,04 | 55,78 ± 5,21 | 1,83 | 36,52 ± 7,10 | 1,2 | 42,87 ± 2,75 | 1,41 |
| | 17-HDHA | 293,26 ± 17,14 | 319,31 ± 32,17 | 1,09 | 335,07 ± 23,99 | 1,14 | 415,17 ± 20,57 | 1,42 | 276,74 ± 54,30 | 0,94 | 369,21 ± 20,83 | 1,26 |
| | 13-HDHA | 109,41 ± 2,84 | 114,21 ± 11,50 | 1,04 | 143,66 ± 13,60 | 1,31 | 156,33 ± 4,32 | 1,43 | 114,27 ± 17,46 | 1,04 | 136,61 ± 4,21 | 1,25 |
| | 10-HDHA | 75,56 ± 1,99 | 80,93 ± 7,61 | 1,07 | 97,51 ± 9,38 | 1,29 | 105,52 ± 3,46 | 1,4 | 79,2 ± 10,41 | 1,05 | 99,51 ± 4,08 | 1,32 |
| | 14-HDHA | 355,14 ± 18,87 | 357,31 ± 22,19 | 1,01 | 328,18 ± 16,86 | 0,92 | 387,62 ± 4,30 | 1,09 | 358,5 ± 46,73 | 1,01 | 415,79 ± 14,74 | 1,17 |
| | 4-HDHA | 789,7 ± 30,37 | 778,07 ± 47,71 | 0,99 | 803,75 ± 42,04 | 1,02 | 868,8 ± 40,91 | 1,1 | 625,13 ± 88,21 | 0,79 | 730,96 ± 35,98 | 0,93 |
| | 13-HODE | 50,21 ± 24,27 | 52,08 ± 26,57 | 1,04 | 353,13 ± 32,15 | 7,03 | 472 ± 33,48 | 9,4 | 87,74 ± 27,48 | 1,75 | 104,66 ± 25,13 | 2,08 |
| | 9-HODE | 38,79 ± 6,83 | 39,39 ± 11,75 | 1,02 | 110,53 ± 18,82 | 2,85 | 115,48 ± 18,77 | 2,98 | 61,6 ± 29,08 | 1,59 | 67,88 ± 17,91 | 1,75 |

**Table S2. Amounts of lipid mediators secreted by human neutrophils.** Concentrations of lipid mediators in the supernatant of untreated and LDC9620-treated human peripheral blood neutrophils after stimulation with *C. albicans* or *S. aureus* as shown in Figure 5A; concentrations are given as pg / 2.25x10[6] neutrophils. Means ± SD of 5 biological replicates per group.

### Supplementary Table S3

| REAGENT | SOURCE | IDENTIFYER |
|---|---|---|
| **Antibodies** | | |
| Armenian hamster monoclonal anti-mouse CD3ε Antibody | Biolegend | Cat #100304; Clone 145-2C11 |
| Donkey polyclonal anti-rabbit IgG | Thermo Fisher Scientific | Cat #A10040 |
| Mouse monoclonal anti-human CD16 Antibody | Biolegend | Cat #302012; Clone 3G8 |
| Mouse monoclonal anti-human CD62L Antibody | Biolegend | Cat #304826; Clone DREG-56 |
| Mouse monoclonal anti-human CD66b Antibody | Biolegend | Cat #305104; Clone G10F5 |
| Mouse monoclonal anti-mouse NK-1.1 Antibody | Biolegend | Cat #108704; Clone PK136 |
| Rabbit monoclonal anti-mouse GLUT1 | Abcam | Cat # ab115730; Clone EPR3915 |
| Rat monoclonal anti-mouse B220 Antibody | Biolegend | Cat #103204; Clone RA-6B2 |
| Rat monoclonal anti-mouse CD115 (CSF-1R) Antibody | Biolegend | Cat #135517; Clone AFS98 |
| Rat monoclonal anti-mouse CD117 (c-Kit) Antibody | Biolegend | Cat # 105814; Clone 2B8 |
| Rat monoclonal anti-mouse CD11b | BioLegend | Cat #101257; Clone M1/70 |
| Rat monoclonal anti-mouse CD16/CD32 | Biozol | Cat #LEIN-C381; Clone 2.4G2 |
| Rat monoclonal anti-mouse CD16/CD32 Antibody | BD Biosciences | Cat #751697; Clone 93 |
| Rat monoclonal anti-mouse CD182 (CXCR2) Antibody | BD Biosciences | Cat #752139; Clone V48-2310 |
| Rat monoclonal anti-mouse CD184 (CXCR4) Antibody | Biolegend | Cat #146511; Clone L276F12 |
| Rat monoclonal anti-mouse CD34 Antibody | Biolegend | Cat # 119314; Clone MEC14.7 |
| Rat monoclonal anti-mouse CD62L | BioLegend | Cat #104412; Clone MEL-14 |
| Rat monoclonal anti-mouse CD90.2 (Thy1.2) Antibody | Biolegend | Cat #140313; Clone 53-2.1 |
| Rat monoclonal anti-mouse Ly-6A/E (Sca-1) Antibody | Biolegend | Cat #108103; Clone D7 |
| Rat monoclonal anti-mouse Ly-6G Antibody | Biolegend | Cat #127608; Clone 1A8 |
| Rat monoclonal anti-mouse Ly-6G/Ly-6C (Gr-1) Antibody | Biolegend | Cat #108427; Clone RB6-8C5 |
| Rat monoclonal anti-mouse Ly6G | BioLegend | Cat #127628; Clone 1A8 |
| Rat monoclonal anti-mouse SiglecF Antibody | BD Biosciences | Cat #740388; Clone E50-2440 |
| Rat monoclonal anti-mouse/human CD11b Antibody | Biolegend | Cat #101243; Clone M1/70 |
| Rat monoclonal anti-mouse/human CD11b Antibody | Biolegend | Cat #101212; Clone M1/70 |
| Rat monoclonal anti-mouse/human CD11b Antibody | Biolegend | Cat #101236; Clone M1/70 |
| Rat monoclonal anti-mouse/human CD11b Antibody | Biolegend | Cat #101208; Clone M1/70 |
| Rat monoclonal anti-mouse/human CD11b Antibody | Biolegend | Cat #101216; Clone M1/70 |
| Streptavidin V500 | BD Biosciences | Cat #561419 |
| **Chemicals and reagents** | | |
| [$^3$H] 2-Deoxy-D-glucose | Perkin Elmer | Cat #NET328A250UC |
| 123count eBeads counting beads | Invitrogen | Cat #01-1234-42 |
| 2-Deoxy-D-glucose (2-DG) | Sigma Aldrich | Cat #D8375 |
| 2-mercaptoethanol | Gibco | Cat #21985023 |
| Antimycin A | Sigma Aldrich | Cat #A8674 |
| Bovine Serum Albumin (BSA) fatty acid free | Sigma Aldrich | Cat #A8806 |
| Cell-Tak | Corning | Cat #354240 |
| D-Glucose | Sigma Aldrich | Cat #G7021 |
| D-Glucose-$^{13}C_6$ | Sigma Aldrich | Cat #389374-1G |
| EDTA | Thermo Fisher Scientific | Cat #15575020 |
| FCS | Sigma Aldrich | Cat #12133 |
| Fixable Viability Dye eFluor 780 | eBioscience | Cat #65-0865-18 |
| GLUT1/3 inhibitor | Lead Discovery Center GmbH | LDC9620 |
| HEPES Buffer Solution | Gibco | Cat #15630-056 |
| IC Fixation Buffer | eBioscience | Cat #00-8222-49 |
| iTaq Universal SYBR Green Supermix | Bio-Rad | Cat #1725124 |
| L-Glutamine | Gibco | Cat #25030081 |
| LB Broth Base | Thermo Fisher Scientific | Cat #12780029 |
| Luminol | Biomol | Cat# Cay16803-5 |
| $NaHCO_3$ | Carl Roth | Cat #27KH.2 |
| NaOH | Carl Roth | Cat #9356.1 |
| Nystatin dihydrate | Carl Roth | Cat #0241.1 |
| Oligomycin | Cayman Chemical | Cat #Cay11341-5 |
| Penicillin-streptomycin | Gibco | Cat #15140122 |

| | | |
|---|---|---|
| Permeabilization Buffer (10X) | eBioscience | Cat#: 00-8333-56 |
| RNAprotect Cell Reagent | Quiagen | Cat #76526 |
| Rotenone | AdipoGen Life Sciences | Cat #AG-CN2-0516-G001 |
| RPMI 1640 GlutaMAX | Gibco | Cat #61870-010 |
| RPMI 1640 w/o glucose | Gibco | Cat #11879020 |
| RPMI 1640 w/o phenol red, w/o glucose | PanBiotech | Cat (#P04-16530 |
| RPMI 1640/o phenol red | Thermofisher | Cat #11835030 |
| Seahorse XF Calibrant Solution, pH 7.4 | Agilent | Cat#: 100840-000 |
| Seahorse XF RPMI medium, pH 7.4 | Agilent | Cat #103576-100 |
| Sodium Pyruvate | Sigma Aldrich | Cat #11360-070 |
| Trifluoromethoxy carbonylcyanide phenylhydrazone (FCCP) | Cayman Chemical | Cat #Cay15218-50 |
| YPD medium | Carl Roth | Cat #X970.2 |
| Zymosan A from *Saccharomyces cerevisiae* | Sigma Aldrich | Cat #4250-1G |
| **Software** | | |
| FACSDiva | BD Bioscience | N/A |
| FlowJo software | FlowJo LLC | https://www.flowjo.com/ |
| GraphPad Prism V10.4.1 | GraphPad Software | https://www.graphpad.com/ |
| Seahorse Wave Software | Agilent | N/A |
| **Critical commercial assays** | | |
| iScript cDNA Synthesis Kit | Bio-Rad | Cat #1708891 |
| MACSxpress® Whole Blood Neutrophil Isolation Kit, human | Miltenyi Biotec | Cat #130-104-434 |
| MS column | Miltenyi Biotec | Cat #130-042-201 |
| Neutrophil Isolation Kit, mouse | Miltenyi Biotec | Cat #130-097-658 |
| PrimeFlow Probe Set Slc2a1 | Thermo Fisher | Cat #VA4-3083505-PF |
| PrimeFlow Probe Set Slc2a3 | Thermo Fisher | Cat #VA4-3083506-PF |
| PrimeFlow™ RNA Assay Kit | Thermo Fisher | Cat #88-18005-204 |
| Roti-Prep RNA Mini | Roth | Cat #8485.2 |
| **Other** | | |
| Aurora Flow Cytometer | Cytek | N/A |
| Celesta Flow Cytometer | BD Bioscience | N/A |
| CFX Connect Real Time System | Bio-Rad | N/A |
| MicroBeta2 microplate scintillation counter | Perkin Elmer | N/A |
| Nanodrop | ThermoFisher | N/A |
| XFe96 Extracellular Flux Analyzer | Seahorse Bioscience | N/A |
| **Primers for qRT-PCR** | | |
| **Target Gene** | **Forward** | **Reverse** |
| Mouse *18S* | CGGCGACGACCCATTCGAAC | GAATCGAACCCTGATTCCCCGT |
| Mouse *Slc2a1* (GLUT1) | GAGACCAAAGCGTGGTGAGT | GCAGTTCGGCTATAACACTGG |
| Mouse *Slc2a3* (GLUT3) | ATCGTGGCATAGATCGGTTC | TCTCAGCAGCTCTCTGGGAT |
| **Experimental models: organisms/strains** | | |
| Mouse: *Mrp8*$^{Cre}$ | Jackson Laboratories | Strain #021614 |
| Mouse: *Slc2a1*$^{fl/fl}$ | Jackson Laboratories | Strain #031871 |
| Mouse: *Slc2a3*$^{fl/fl}$ | Provided by E. Dale Abel (Iowa) | Fidler et al., 2017 |
| Mouse: C57BL/6 | Jackson Laboratories | Strain #000664 |
| *Staphylococcus aureus* (HG001) | Provided by K. Ohlsen (JMU Würzburg) | Herbert et al., 2010 |
| *Candida albicans* (SC5314) | Provided by J. Morschhäuser (JMU Würzburg) | Gillum et al., 1984 ATCC: MYA-2876 |

**Table S3. Reagents used in this study**
